# Actin crosslinking is required for force sensing at tricellular junctions

**DOI:** 10.1101/2025.02.21.639590

**Authors:** Nilay Taneja, Michael F. Moubarak, Meriel J. McGovern, Kenji Yeoh, Jennifer A. Zallen

## Abstract

Mechanical forces are essential for tissue morphogenesis, but risk causing ruptures that could compromise tissue function. In epithelial tissues, adherens junctions withstand the forces that drive morphogenesis by recruiting proteins that stabilize cell adhesion and reinforce connections to the actin cytoskeleton under tension. However, how junctional actin networks respond to forces *in vivo* is not well understood. Here we show that the actin crosslinker Fimbrin is recruited to tricellular junctions under tension and plays a central role in amplifying actomyosin contractility and stabilizing cell adhesion. Loss of Fimbrin results in a failure to reorganize actin under tension and an inability to enhance myosin-II activity and recruit junction-stabilizing proteins in response to force, disrupting cell adhesion. Conversely, increasing Fimbrin activity constitutively activates force-response pathways, aberrantly stabilizing adhesion. These results demonstrate that Fimbrin-mediated actin crosslinking is an essential step in modulating actomyosin dynamics and reinforcing cell adhesion under tension during epithelial remodeling.

## Introduction

The organization of the actin cytoskeleton into force-generating networks is necessary for the dynamic cell behaviors that build multicellular tissues (*1*). Cells in epithelial sheets are exposed to mechanical forces that are sensed by load-bearing cell-cell and cell-matrix adhesions and activate force-regulated pathways that influence cell fate, cell behavior, and tissue structure (*2-4*). In particular, tricellular junctions are sites of increased forces that control cell shape, differentiation, division, and barrier function (*5, 6*). These forces are essential drivers of morphogenesis, but require mechanisms to prevent breaks or tears that could disrupt tissue integrity and function. During epithelial remodeling, cells maintain tissue integrity by recruiting proteins to tricellular junctions that stabilize cell adhesion under tension, including the core adherens junction protein E-cadherin (*7, 8*) and the junction-actin linker proteins Vinculin (*9*), Canoe/Afadin (*10-12*), and Ajuba (*13*). In addition, *in vitro* studies show that mechanical forces can stabilize interactions between adhesion receptors (*14, 15*), between adhesion complex components (*16-18*), and between junctional proteins and the actin cytoskeleton (*19-21*). However, despite the essential role of the actin cytoskeleton in supporting cell adhesion (*3, 22*), the actin cytoskeleton is a highly dynamic structure that turns over on the order of seconds (*23*) and rapidly organizes into diverse structures through the activities of actin nucleating, crosslinking, and motor proteins (*24-26*). How adherens junctions stably anchor to dynamic cytoskeletal networks is not well understood. Several actin regulators preferentially interact with actin filaments under tension *in vitro* (*27-34*) and mechanical forces can modulate actin polymerization (*35-37*), organization (*38*), and repair (*39, 40*) in cultured cells. However, although actin is enriched at tricellular junctions (*38, 41*), how junctional actin networks sense and respond to physiological forces *in vivo*, and whether force-induced changes in the actin cytoskeleton are important for cell adhesion, is not known.

## Results

### F-actin organization at epithelial tricellular junctions is regulated by contractile forces

To investigate the effects of mechanical forces on actin organization, we used Airyscan imaging (*42*) and deconvolution to visualize the localization of filamentous actin (F-actin) in the *Drosophila* embryo (Figs. 1A and S1A), focusing on tricellular junctions that are exposed to dynamic forces during epithelial remodeling (*43*). Based on the estimated thickness of cortical actin networks in cultured cells (*44*), we analyzed cortical F-actin organization within 0.5 μm of the tricellular junction by measuring the ratio of F-actin signal in the periphery of this region to the F-actin signal closer to the membrane (Figs. 1B and 1C). This measurement, referred to as the F-actin span, varied over a nearly 10-fold range (Fig. 1C), indicating that F-actin is more concentrated at the membrane in some tricellular junctions and more diffusely distributed in others.

**Figure 1.**
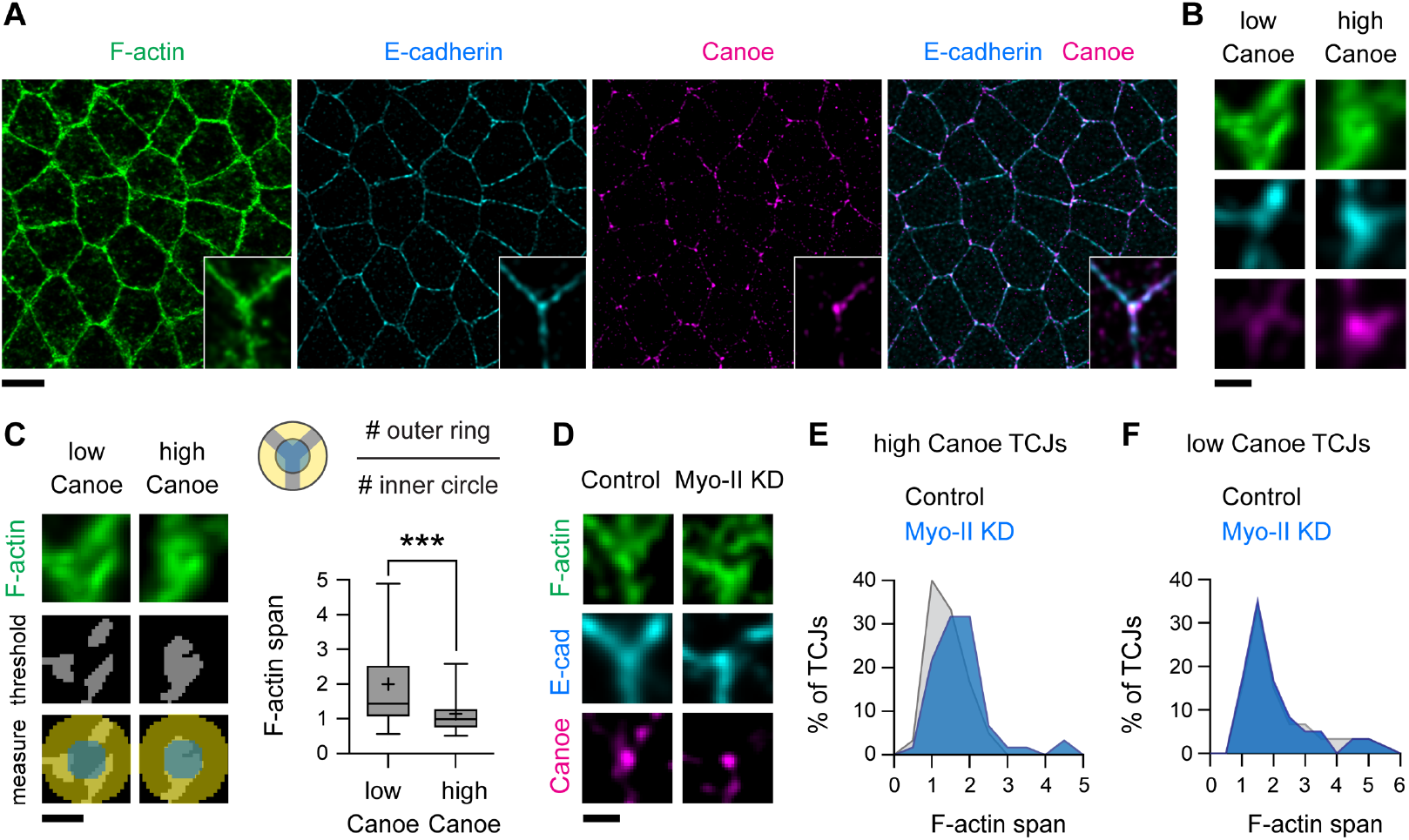
Actin reorganization at tricellular junctions under tension. **(A)** Localization of F-actin (phalloidin), E-cadherin, and Canoe in wild-type (WT) embryos. Insets, close-ups of single tricellular junctions (TCJs). **(B)** Examples of low Canoe and high Canoe TCJs. (**C)** F-actin span values at TCJs (# positive pixels in the outer ring divided by # positive pixels in the inner circle after thresholding) were significantly different at low Canoe (bottom quartile) and high Canoe (top quartile) TCJs in WT embryos. **(D)** Examples of high Canoe TCJs in control (Gal4 only) and Myo-II KD embryos. **(E, F)** F-actin span values were significantly different at high Canoe TCJs in control and Myo-II KD embryos (p<0.05, unpaired t-test with Welch’s correction, see Fig. S1G). No difference was detected at low Canoe TCJs (see Fig. S1H). Fixed stage 7-8 embryos are shown in all panels, oriented ventral down in (A), 220-240 TCJs in 5-6 embryos/genotype. Boxes, 25^th^-75^th^ percentile; whiskers, 5^th^-95^th^ percentile; horizontal line, median; +, mean. ***, p<0.001, unpaired t-test with Welch’s correction. Bars, 5 μm in (A) and 0.5 μm in (B-D).

To determine if these differences in actin organization are due to variations in force, we used the actin-junction linker protein Canoe as a readout of mechanical forces in the embryo, as Canoe is recruited to tricellular junctions in response to actomyosin contractility (*10*) (Figs. S1B-S1E). Tricellular junctions with the highest levels of Canoe (the top 25% of tricellular junctions) had a more focused concentration of F-actin at the membrane, whereas tricellular junctions with the lowest levels of Canoe (the bottom 25%) displayed a more dispersed distribution (Figs. 1B and 1C). These results demonstrate that F-actin organization correlates with a marker of tension, with total F-actin levels remaining constant (Fig. S1F).

To test if mechanical forces are necessary for actin organization, we expressed a short hairpin RNA targeting the myosin-II heavy chain, which reduces cortical tension in the *Drosophila* embryo by ∼50% (*45*). Canoe levels at tricellular junctions were significantly reduced in *myosin-II* KD embryos (Figs. S1D and S1E) consistent with an overall decrease in myosin activity, although Canoe levels still varied over a broad range, likely due to local variations in residual tension. In *myosin-II* KD embryos, the actin span was increased at tricellular junctions with the highest Canoe levels, but was unchanged at low-Canoe junctions, with no difference in F-actin levels (Figs. 1D-1F and S1G-S1I). These results demonstrate that actin organization at tricellular junctions requires myosin-II activity, consistent with the regulation of these actin structures by mechanical force.

### The actin-crosslinking protein Fimbrin is recruited to tricellular junctions under tension

To investigate how mechanical forces influence actin organization, we looked for actin regulators that localize to tricellular junctions under tension. Through a screen of candidate actin-binding proteins (Figs. S2A and S2B), we found that the actin crosslinker Fimbrin, a conserved member of the Fimbrin/Plastin protein family (*46, 47*), was enriched 2.5±0.9-fold (mean±SD) at tricellular junctions in the *Drosophila* embryo (Figs. 2A-2C). A Fimbrin-YFP fusion expressed from the endogenous locus (*48*) colocalized with myosin-II at tricellular and bicellular junctions (Figs. S2C-S2H) and correlated with changes in myosin-II levels over time (Figs. 2D and S2I). However, despite essential roles for Fimbrin in several processes that require mechanical force, such as cytokinesis, endocytosis, and cell migration (*49-55*), whether Fimbrin is required for cells to respond to force is unknown.

**Figure 2.**
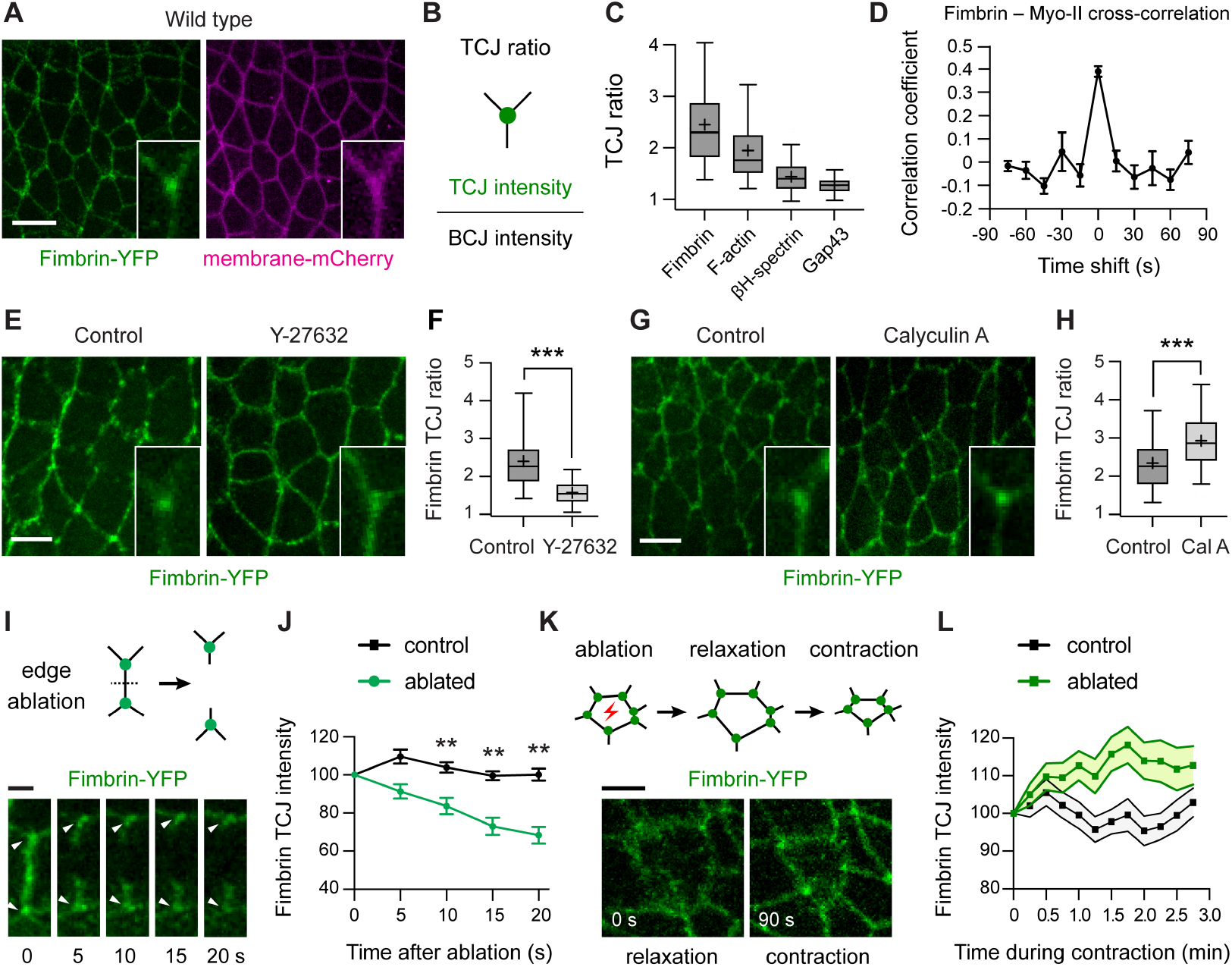
The Fimbrin actin-crosslinking protein is recruited to tricellular junctions under tension. **(A)** Localization of Fimbrin-YFP in wild-type (WT) embryos. Gap43-mCherry, membrane marker. **(B, C)** TCJ ratio (TCJ intensity divided by the mean intensity of the three connected BCJs) of Fimbrin-YFP, F-actin (Moesin actin-binding domain-mCherry), the related crosslinker βH-spectrin-YFP, and the membrane marker Gap43-mCherry in WT embryos. **(D)** Temporal correlation between Fimbrin-YFP and myosin-II-mCherry at TCJs. **(E, F)** Localization (E) and TCJ ratio (F) of Fimbrin-YFP in control (water-injected) and Y27632-injected embryos. **(G, H)** Localization (G) and TCJ ratio (H) of Fimbrin-YFP in control (DMSO-injected) and Calyculin A-injected embryos. Insets, close-ups of single TCJs. **(I, J)** Fimbrin-YFP localization (I) and intensity (J) at TCJs after tension release by edge ablation, normalized to 100% prior to ablation (t=0). Arrowheads, TCJs before (0 s) and after (5-20 s) edge ablation. **(K, L)** Fimbrin-YFP localization (K) and intensity (L) during the contraction phase following apical ablation, normalized to 100% at t=0, the time of maximal relaxation. Live stage 7-8 embryos are shown in all panels, oriented ventral down in (A), (E), and (G), 121-220 TCJs in 4-9 embryos/genotype in (C), (F), and (H), 22-36 TCJs/condition in (D), (J), and (L). Mean±SEM in (D), (J), and (L). Boxes, 25^th^-75^th^ percentile; whiskers, 5^th^-95^th^ percentile; horizontal line, median; +, mean. **, p<0.001, ***, p<0.0001 (unpaired t-test with Welch’s correction). Bars, 10 μm in (A), 5 μm in (E), (G), and (K) and 2 μm in (I).

If Fimbrin is important for the cellular response to force, then its localization or activity is predicted to be regulated by actomyosin contractility, the predominant driver of mechanical forces in the early *Drosophila* embryo (*43*). To investigate whether actomyosin forces influence Fimbrin localization, we analyzed the effects of pharmacological inhibitors that alter myosin-II activity. Injection of embryos with the Rho-kinase inhibitor Y-27632, which decreases myosin-II activity (*56*), reduced Fimbrin-YFP enrichment at tricellular junctions by 57% (Figs. 2E and 2F). By contrast, injection of the myosin-II phosphatase inhibitor Calyculin A, which increases myosin-II activity (*57*), enhanced Fimbrin-YFP enrichment at tricellular junctions by 43% (Figs. 2G and 2H). These results demonstrate that Fimbrin localization at tricellular junctions is regulated by myosin-II activity.

To directly test whether Fimbrin localization is regulated by force, we used laser ablation to manipulate mechanical forces at tricellular junctions. Ablation of single cell edges, which rapidly reduces tension at the associated tricellular junctions (*10*), decreased Fimbrin-YFP intensity at tricellular junctions by nearly 30% (Figs. 2I and 2J). Edge ablation did not significantly affect the localization of another actin-crosslinking protein, βH-spectrin, indicating that Fimbrin dissociation is not due to a general loss of F-actin (Fig. S2J). By contrast, laser ablation of the apical cell cortex, which locally enhances tension at tricellular junctions by inducing contraction of the ablated cell (*12, 58*), increased Fimbrin-YFP intensity at tricellular junctions by nearly 20% (Figs. 2K, 2L, and Movie S1). These results demonstrate that mechanical forces are necessary and sufficient for Fimbrin localization to tricellular junctions.

### Fimbrin is necessary for force-regulated changes in actin organization and cell adhesion

The recruitment of Fimbrin to tricellular junctions under tension raised the possibility that Fimbrin could be involved in mediating force responses at these structures. To test this, we used CRISPR/Cas9-mediated genome engineering to generate a null allele that lacks most of the *Fimbrin* open reading frame (Fig. S3A). The *Fimbrin* null allele significantly reduced embryo viability and this defect was rescued by restoring the wild-type gene, indicating that this mutation is specific (Fig. S3B). The average F-actin span at tricellular junctions in *Fimbrin* mutants was increased by more than 30% compared to wild-type embryos (Figs. 3A-3C), reminiscent of tricellular junctions under low tension in wild type, with no change in total F-actin levels (Fig. S3C). This increase in F-actin span is consistent with a role for Fimbrin in generating or responding to force. To test if Fimbrin is required to generate force, we analyzed myosin-II localization and activity in *Fimbrin* mutants. Myosin-II localized correctly to cell edges in a planar polarized fashion in *Fimbrin* mutants, similar to wild-type embryos (Figs. S3D and S3E) (*59, 60*). To analyze myosin-II contractility, we used laser ablation to sever single cell edges, as the peak retraction velocity after ablation is predicted to correlate with the tension on the edge prior to ablation (*61, 62*). Single cell edges in *Fimbrin* mutants generated wild-type forces measured by laser ablation, indicating that baseline myosin-II activity occurs normally in the absence of Fimbrin (Fig. S3F). These results demonstrate that Fimbrin is not required to generate myosin-II contractility, but is necessary for actin reorganization in response to tension at tricellular junctions.

**Figure 3.**
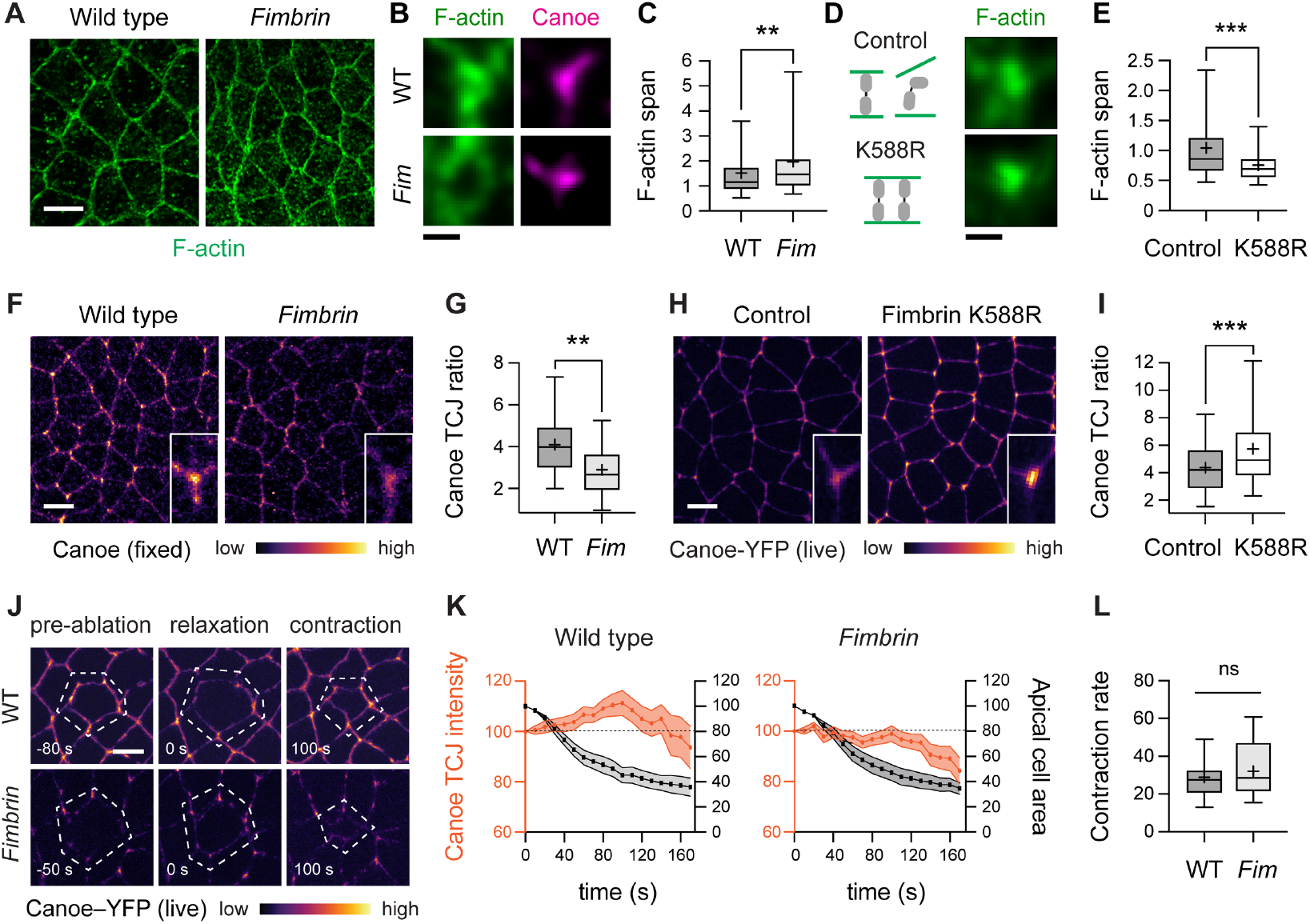
Fimbrin is necessary and sufficient to activate force responses at tricellular junctions. **(A**) Localization of F-actin (phalloidin) in wild-type (WT) and *Fimbrin* mutant (*Fim*) embryos. (**B**) Canoe and F-actin localization in examples of high Canoe TCJs (top quartile) in WT and *Fim* embryos. (**C**) F-actin span values at all TCJs in WT and *Fim* embryos. **(D)** Localization of F-actin (Utrophin-mCherry) at TCJs in control (Gal4 only) and Fimbrin^K588R^ embryos. **(E)** F-actin span values at all TCJs in control and Fimbrin^K588R^ embryos. (**F, G**) Canoe localization (F) and TCJ ratio (G) in WT and *Fim* embryos. **(H, I)** Canoe-YFP localization (H) and TCJ ratio (I) in control and Fimbrin^K588R^ embryos. Insets, close-ups of single TCJs. **(J)** Canoe-YFP localization before apical ablation, at maximal relaxation (t=0 s), and during the contraction phase after apical ablation. Dotted outline, ablated cell. **(K)** Canoe-YFP TCJ intensity (orange) and apical area of the ablated cell (black) during the contraction phase in WT and *Fim* embryos (p<0.05 at t=100 s, unpaired t-test with Welch’s correction). Measurements were normalized to 100% at t=0, the time of maximal relaxation. **(L)** Apical area contraction rates (% area decrease/min) were similar in WT (29±3%/min) and *Fim* (32±4%/min) embryos. Fixed stage 7-8 embryos are shown in (A-C), (F), and (G) and live stage 7-8 embryos are shown in (D), (E), and (H-L), oriented ventral down in (A), (F), and (H), 176-240 TCJs in 5-7 embryos/genotype in (C), (E), and (I), 309-367 TCJs in 7-8 embryos/genotype in (G), and 42 TCJs from 11 cells in 10-11 embryos/genotype in (K) and (L). Mean±SEM in (K). Boxes, 25^th^-75^th^ percentile; whiskers, 5^th^-95^th^ percentile; horizontal line, median; +, mean. ns, not significant. **, p<0.001, ***, p<0.0001 (unpaired t-test with Welch’s correction). Bars, 5 μm in (A), (F), (H), and (J), 500 nm in (B) and (D).

The effect of Fimbrin on actin organization raised the question of whether Fimbrin is required for other force responses at tricellular junctions. Mechanical forces recruit several proteins to tricellular junctions, including the adherens junction protein E-cadherin (*7, 8*) and the junction-actin linker proteins Canoe (*10-12*), and Ajuba (*13*), which reinforce cell adhesion under tension. To investigate whether Fimbrin is required to recruit junction-stabilizing proteins, or alternatively, if force-induced changes in junctional composition are upstream or independent of Fimbrin, we analyzed protein localization in wild-type and *Fimbrin* mutant embryos. Canoe enrichment at tricellular junctions was reduced by nearly 40% in *Fimbrin* mutants (Figs. 3F and 3G), whereas Fimbrin localization in *canoe* KD embryos occurred normally (Fig. S4A). *Fimbrin* mutants also displayed a significant reduction in the enrichment of E-cadherin and Ajuba at tricellular junctions (Figs. S4B and S4C). These results demonstrate that Fimbrin is required for the localization of Canoe and other force-sensitive proteins to tricellular junctions.

As Canoe is regulated by force-dependent (*10-12*) and force-independent (*63-66*) inputs, we tested if Fimbrin is required for Canoe localization under tension. Canoe-GFP is recruited to tricellular junctions by ectopic forces induced by apical ablation (*12*). Canoe recruitment in response to apical ablation was abolished in *Fimbrin* mutants (Figs. 3J, 3K, Movie S2, and Movie S3). Despite a failure to recruit Canoe, apical contraction occurred normally, indicating that Fimbrin is not required to generate ectopic forces (Fig. 3L). Moreover, Canoe localization defects in *Fimbrin* mutants were first detected in stage 7, when Canoe localization is force-dependent, and Fimbrin was dispensable for Canoe localization in stage 6, when Canoe is regulated by force-independent signals (Figs. 3F, 3G, and S4D). Similar defects were observed in embryos in which Fimbrin protein was depleted by nanobody-mediated protein degradation (referred to as Fim KD embryos) (Figs. S4E-S4H). Together, these results indicate that Fimbrin is specifically required to recruit Canoe to tricellular junctions under tension.

### Increased Fimbrin activity recapitulates the effects of forces at tricellular junctions

As Fimbrin regulates actin organization at tricellular junctions (Figs. 3A-3C), and actin is required for cell adhesion (*3, 22*), it is perhaps not surprising that the loss of Fimbrin disrupts protein localization at adherens junctions. However, if Fimbrin is an instructive regulator of force responses, then Fimbrin should not only be necessary, but also sufficient to induce force responses at tricellular junctions. To test this possibility, we engineered a Fimbrin gain-of-function mutation identified in *S. cerevisiae* (Fimbrin^K610R^) (*67*) into the corresponding location in a *Drosophila* Fimbrin transgene (Fimbrin^K588R^). Fimbrin contains two actin-binding domains that are both required for Fimbrin localization to tricellular junctions (Figs. S5A-S5C). The Fimbrin^K588R^ mutation, located in the second actin-binding domain (Fig. S5C), is predicted to enhance Fimbrin binding to actin based on the cryo-EM structure of mammalian Plastin (*68*). Consistent with increased actin association, Fimbrin^K588R^ was more stably associated with the cortex in *Drosophila* S2R+ cells, resulting in decreased fluorescence recovery after photobleaching, a measure of protein turnover at the cell cortex (Figs. S5D-S5G). Moreover, Fimbrin^K588R^ localized to tricellular junctions to nearly the same extent as wild-type Fimbrin (Figs. S5A and S5B), providing an opportunity to examine how this variant affects force responses at tricellular junctions. Maternal expression of Fimbrin^K588R^ reduced the F-actin span at tricellular junctions by 25% (Figs. 3D and 3E) and enhanced the enrichment of Canoe by 38% (Figs. 3H and 3I), two properties of tricellular junctions under tension. These results demonstrate that Fimbrin^K588R^ is sufficient to mimic the effects of forces on protein localization and cytoskeletal organization at tricellular junctions.

### Fimbrin modulates the rate of cell rearrangement during axis elongation

The findings that Fimbrin is required for actin organization and Canoe localization at tricellular junctions under tension, and can reproduce the effects of forces when activated, suggest that Fimbrin plays an essential role in converting mechanical forces into changes in cell behavior. To investigate this possibility, we analyzed the effects of Fimbrin on cell adhesion and epithelial remodeling, two cellular outputs of forces in the *Drosophila* embryo. In particular, the force-regulated recruitment of Canoe to tricellular junctions is required to stabilize cell adhesion and tune the rate of cell rearrangement under tension (*10-12*). Cell adhesion is normally maintained throughout axis elongation, with gaps in E-cadherin localization occurring rarely, at 3±1% (mean±SEM) of 3-cell junctions and 7±2% of 4-cell junctions that encounter strong forces during cell rearrangement (Figs. 4A-4C). By contrast, E-cadherin gaps occurred at 21±2% of 3-cell junctions and 35±3% of 4-cell junctions in *Fimbrin* mutants (Figs. 4A-4C). E-cadherin defects were first observed in stage 7 when mechanical forces are upregulated at the onset of axis elongation and were not detected prior to elongation in stage 6 (Figs. S6A and S6B). Fim KD embryos displayed similar defects (Figs. S6C-S6G). These results demonstrate that Fimbrin is necessary to maintain adhesion at tricellular and 4-cell junctions during axis elongation.

**Figure 4.**
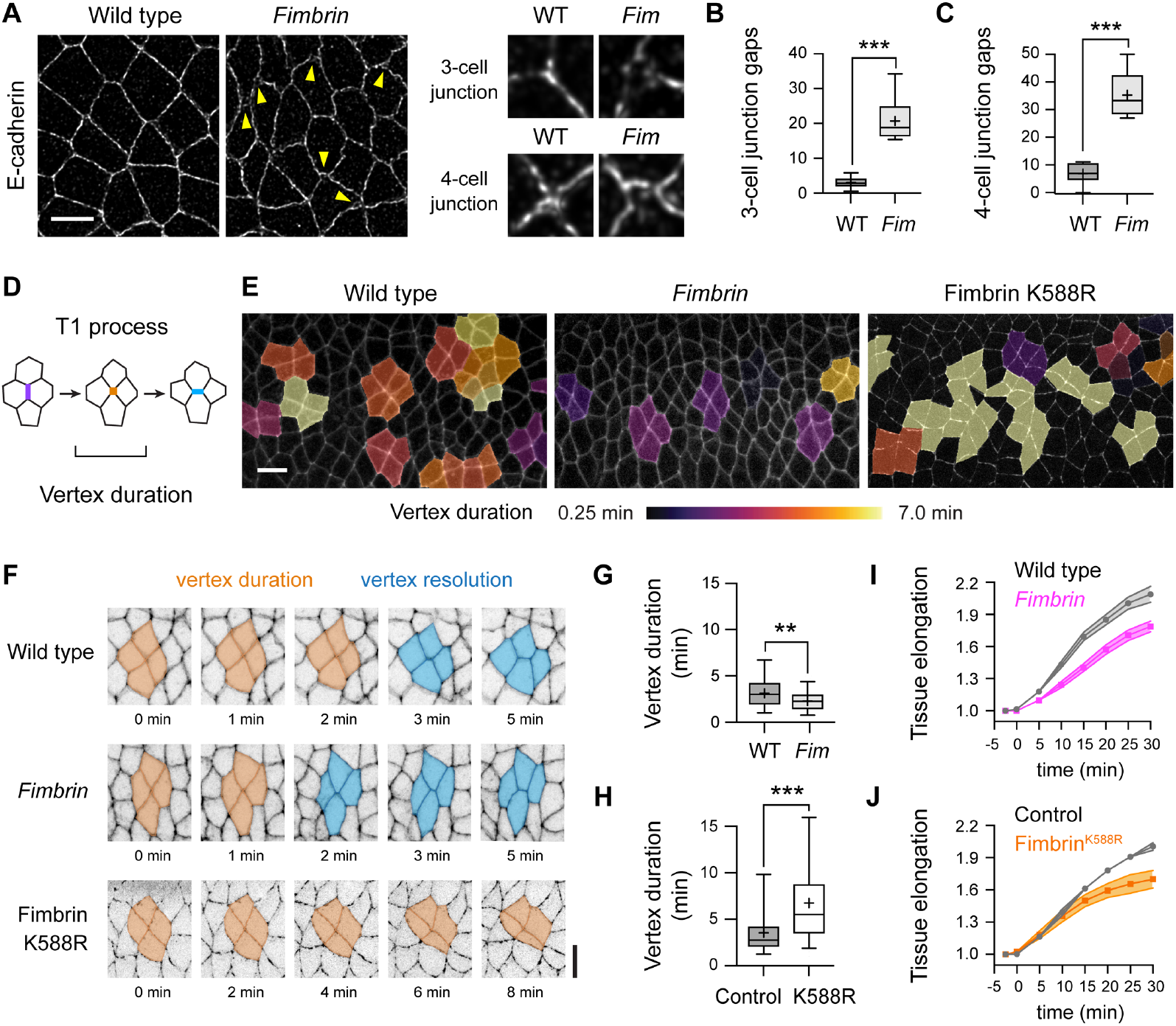
Fimbrin regulates cell adhesion and cell rearrangement during axis elongation. **(A)** E-cadherin localization in wild-type (WT) and *Fimbrin* mutant (*Fim*) embryos. Arrowheads, junctional gaps. **(B, C)** Percentage of 3-cell and 4-cell junctions with gaps in WT and *Fim* embryos. **(D)** Cell rearrangement (T1 process) schematic. An edge oriented parallel to the dorsal-ventral axis (vertical edge, purple) contracts to form a 4-cell junction (orange) that resolves through the formation of a new edge perpendicular to this axis (blue). **(E)** Stills from movies of WT, *Fim*, and Fimbrin^K588R^ embryos expressing Spider-GFP (WT and *Fim*) or -catenin-GFP (Fimbrin^K588R^) 12.5 min after the start of elongation showing cells in T1 processes color-coded by vertex duration. **(F)** Vertex resolution in WT, *Fim*, and Fimbrin^K588R^ embryos. Orange, cells meet at a 4-cell vertex. Blue, cells form a new contact. **(G, H)** Duration of 4-cell junctions (vertex duration) in WT and *Fim* embryos (G) and control (Gal4 only) and Fimbrin^K588R^ embryos (H). **(I, J)** Tissue elongation (tissue length normalized to the length at t=0) in WT and *Fim* (I), and control and Fimbrin^K588R^ (J) embryos. p<0.05 (unpaired t-test with Welch’s correction) at t=30 min for *Fimbrin* and Fimbrin^K588R^. Fixed stage 7-8 embryos are shown in (A-C), live stage 7-8 embryos are shown in (E-J), oriented ventral down in (A), (E), and (F), 1,297-1,322 3-cell junctions and 161-187 4-cell junctions in 8-10 embryos/genotype in (B) and (C), 81-90 4-cell junctions in 3-4 embryos/genotype in (G) and (H), and 3-4 embryos/genotype in (I) and (J). Boxes, 25^th^-75^th^ percentile; whiskers, 5^th^-95^th^ percentile; horizontal line, median; +, mean. **, p<0.01. ***, p<0.0001 (unpaired t-test with Welch’s correction). Bars, 5 μm in (A) and (F), 10 μm in (E).

To test if Fimbrin is required for epithelial remodeling, we generated time-lapse movies of axis elongation in embryos with increased or decreased Fimbrin activity (Movies S4 and S5), using computational tools for image segmentation and analysis (Materials and Methods) (*69*). In wild-type embryos, tissue elongation along the head-to-tail axis is driven by spatially regulated cell rearrangements in which edges oriented perpendicular to the head-to-tail axis (vertical edges) contract to form junctions where four or more cells meet, and these junctions resolve by forming new cell contacts (Fig. 4D) (*59, 70*). If Fimbrin stabilizes cell adhesion under tension, then loss of Fimbrin is predicted to destabilize adhesion and accelerate rearrangement, whereas increased Fimbrin activity is predicted to aberrantly stabilize adhesion, causing cell rearrangements to slow down or stall. Nearly all vertical edges contracted in *Fimbrin* mutant and Fimbrin^K588R^ embryos, similar to wild type, indicating that altering Fimbrin activity does not prevent cell rearrangement (Fig. S6H). By contrast, 4-cell junctions resolved faster in the absence of Fimbrin, with 4-cell junctions resolving within 2.3±1.1 min (mean±SD) in *Fimbrin* mutants compared with 3.1±1.7 min in wild type (Figs. 4E-4H). These defects were accompanied by the formation of aberrant epithelial folds, consistent with a failure to maintain tissue integrity (Fig. S6I). Conversely, 4-cell junctions resolved more slowly in Fimbrin^K588R^ embryos, remaining stable for nearly twice as long as in controls (Figs. 4E, 4F, and 4H). These results demonstrate that increasing or decreasing Fimbrin activity have opposite effects on junctional remodeling, suggesting that Fimbrin activity tunes the rate of cell rearrangement. Both *Fimbrin* mutant and Fimbrin^K588R^ embryos displayed fewer total cell rearrangements overall (Fig. S7) and a reduction in axis elongation (Figs. 4I and 4J), indicating that an optimal level of Fimbrin activity is necessary for proper epithelial morphogenesis.

### Fimbrin amplifies myosin-II contractility under tension

The effects of Fimbrin on actin organization, protein localization, and cell rearrangement indicate that Fimbrin is necessary and sufficient for multiple force responses at tricellular junctions. How Fimbrin carries out these diverse functions is not known. In one model, Fimbrin could generate an actin structure that provides a stable substrate for the recruitment of actin-junction linker proteins that reinforce the connection between adherens junctions and the cytoskeleton. Alternatively, Fimbrin could act at a distinct step to promote the recruitment of myosin-II, which could create a positive feedback loop that amplifies forces at tricellular junctions and activates multiple tension-sensitive pathways. The amplification of myosin-II activity by mechanical feedback shapes the distribution of forces in several cell and tissue contexts. Mechanical regulation of myosin-II contractility in epithelia results in the formation of multicellular contractile cables that drive cell rearrangement, establish compartment boundaries, and maintain tissue structure under stress (*70-78*). However, whether mechanical regulation of myosin-II contributes to the force response at tricellular junctions, and if Fimbrin-dependent actin crosslinking regulates myosin-II activity under force, is unknown.

To address these questions, we analyzed myosin-II localization and dynamics at tricellular junctions in *Fimbrin* mutant and Fimbrin^K588R^ embryos. Three lines of evidence indicate that Fimbrin regulates myosin-II localization and dynamics at tricellular junctions under tension. First, myosin-II levels at tricellular junctions were significantly decreased in *Fimbrin* mutants and increased in Fimbrin^K588R^ embryos, demonstrating that Fimbrin is necessary and sufficient for myosin-II localization (Figs. 5A-5D). Second, fluorescence recovery after photobleaching experiments showed that loss of Fimbrin enhances myosin-II-GFP turnover, measured as an increase in the fraction of myosin-II-GFP fluorescence recovered 35 s after bleaching the tricellular junction (the mobile fraction) (Figs. 5E and 5F). Conversely, Fimbrin^K588R^ expression inhibited myosin-II-GFP turnover at tricellular junctions, decreasing the mobile fraction (Figs. 5E and 5F). These results indicate that Fimbrin is necessary and sufficient to stabilize myosin-II cortical localization at tricellular junctions. In a third approach, we found that myosin-II levels at tricellular junctions were increased by more than 30% in response to ectopic forces induced by apical ablation, indicating that myosin-II localization is tension-sensitive (Figs. 5I, 5J, and Movie S6).This recruitment was eliminated in *Fimbrin* mutants, demonstrating that Fimbrin is required for myosin-II recruitment in response to tension (Figs. 5I, 5J, Movie S6, and Movie S7). Together, these results demonstrate that Fimbrin is necessary and sufficient to stabilize myosin-II localization in response to mechanical forces at tricellular junctions.

**Figure 5.**
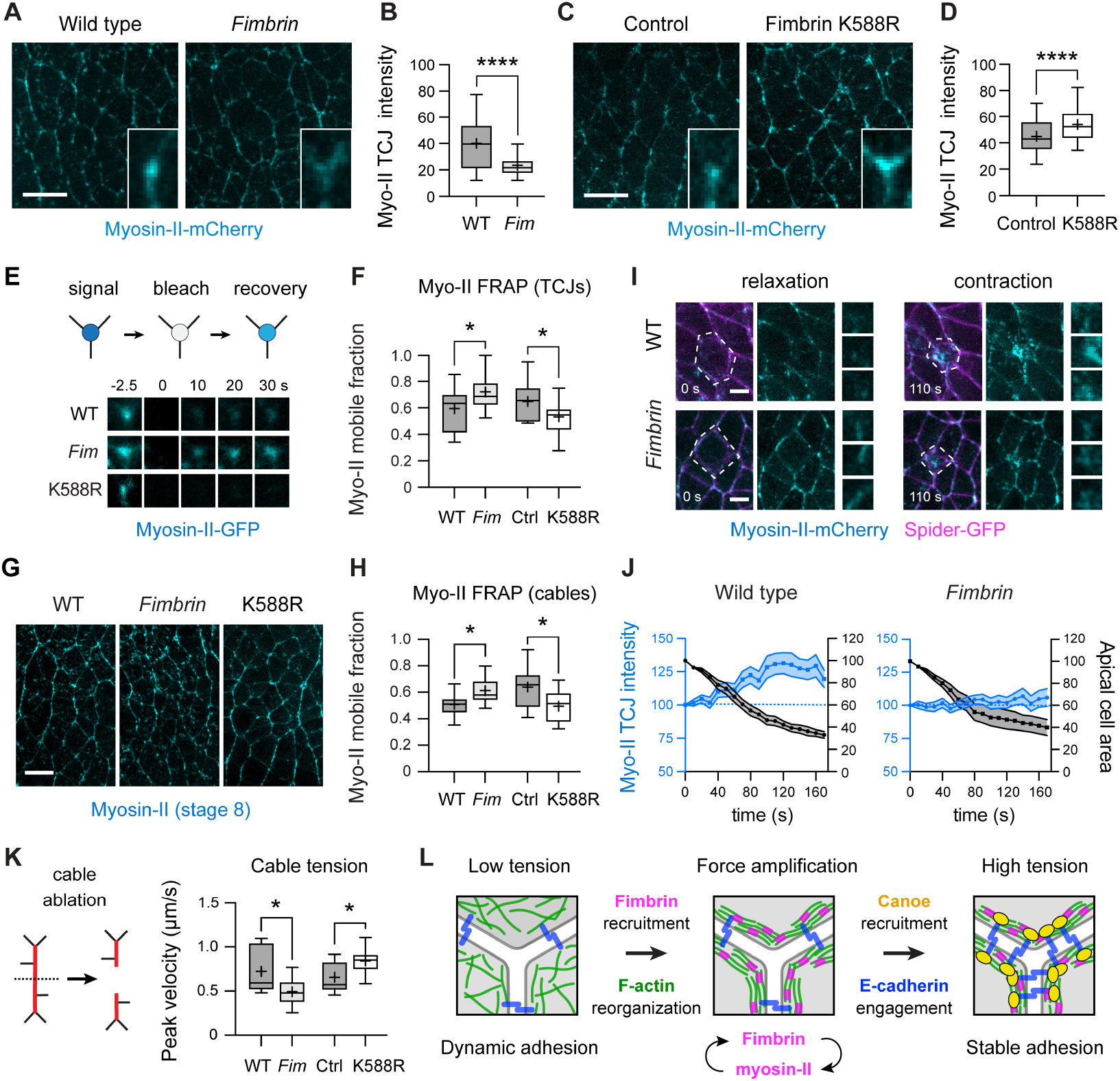
Fimbrin modulates myosin-II dynamics in multiple cortical domains under tension. **(A-D)** Myosin-II-mCherry localization (A, C) and intensity (B, D) in wild-type (WT) and *Fimbrin* mutant (*Fim*) embryos (A, B) and control (Gal4 only) and Fimbrin^K588R^ embryos (C,D). **(E, F)** Myosin-II-GFP fluorescence recovery after photobleaching (E) and mobile fraction (F) at TCJs in WT, *Fim*, control, and Fimbrin^K588R^ embryos. **(G, H)** Localization (G) and mobile fraction (H) of myosin-II in multicellular cables (≥3 connected edges oriented within 30° of the dorsal-ventral axis) in WT and *Fim* (expressing Myosin-II-GFP), and control and Fimbrin^K588R^ (expressing Myosin-II-mCherry). **(I)** Myosin-II-mCherry localization at maximal relaxation (0 s) and during the contraction phase following apical ablation (110 s). Spider-GFP, membrane marker. Dotted outline, ablated cell. Crops, examples of individual TCJs. **(J)** Myosin-II-mCherry TCJ intensity (blue) and apical area of the ablated cell (black) during the contraction phase in WT and *Fim* embryos (p<0.05 at t=100 s, unpaired t-test with Welch’s correction). Measurements were normalized to 100% at t=0, the time of maximal relaxation. Apical area contraction rates (% area decrease/min) after ablation were similar in WT (23±1%/min) and *Fim* (21±2%/min). **(K)** Ablation of myosin-II cables (schematic, left) and peak velocity measurements in WT, *Fim*, control, and Fimbrin^K588R^ embryos. **(L)** Fimbrin (magenta) promotes actin reorganization, force amplification, and cell adhesion through the recruitment of junction-stabilizing proteins. Blue, E-cadherin. Live stage 7-8 embryos are shown in all panels, oriented ventral down, 169-237 TCJs in 4-5 embryos/genotype in (B) and (D), 12-32 TCJs/genotype in (F) and (J), 9-17 cables/genotype in (H) and (K). Mean±SEM in (J). Boxes, 25^th^-75^th^ percentile; whiskers, 5^th^-95^th^ percentile; horizontal line, median; +, mean. *, p<0.05, ****, p<0.0001 (unpaired t-test with Welch’s correction). Bars, 10 μm in (A), (C) and (G), and 5 μm in (I).

The effects of Fimbrin on myosin-II localization are consistent with a model in which Fimbrin is required to amplify actomyosin contractility in response to mechanical forces at tricellular junctions. As Fimbrin is also present at bicellular junctions (Figs. 2A, S2G, and S2I), this raised the question of whether Fimbrin also influences force responses in other cellular domains. Myosin-II localization is initiated at single edges in the *Drosophila* embryo by biochemical cues (*79*), and subsequently amplified by mechanical feedback to promote the assembly of multicellular myosin-II cables, which generate increased contractile forces that drive cell rearrangement (*70, 71*). If Fimbrin is specifically required for mechanical feedback, then perturbing Fimbrin should affect the amplification but not the initiation of myosin-II contractility. To examine whether Fimbrin is required for mechanical feedback, we compared the effects of Fimbrin on myosin-II behavior in cables and single edges. We found that myosin-II localization, dynamics, and activity at single vertical edges were unaffected in *Fimbrin* mutant, Fim KD, and Fimbrin^K588R^ embryos, indicating that increasing or decreasing Fimbrin does not affect myosin-II regulation at single edges (Figs. S3D-S3F and S8A-S8G). By contrast, modifying Fimbrin activity had striking effects on myosin-II cables. Fewer cables formed in *Fimbrin* mutant and Fim KD embryos, and the cables that were present were shorter and displayed increased myosin-II turnover compared to wild-type embryos (Figs. 5G, 5H, and S8H-S8N). Conversely, myosin-II cables were longer on average in Fimbrin^K588R^ embryos and displayed decreased myosin-II turnover, indicating that Fimbrin stabilizes myosin-II cortical localization in these structures (Figs. 5G, 5H, and S8M). These effects were associated with significant changes in myosin-II contractility. Contractile forces generated by myosin-II cables were decreased in *Fimbrin* mutant and Fim KD embryos and increased in Fimbrin^K588R^ embryos, whereas contractile forces at isolated vertical edges were unaffected (Figs. 5K, S3F, and S8O-S8Q). Collectively, these results demonstrate that Fimbrin regulates myosin-II localization, dynamics, and activity at tricellular junctions and in multicellular myosin-II cables, indicating that Fimbrin participates in a broadly acting mechanism required to amplify mechanical forces under tension in the *Drosophila* embryonic epithelium.

## Discussion

Cells maintain tissue integrity during epithelial remodeling through force-activated pathways that modulate cell adhesion under tension, but how these molecular responses are integrated with the cytoskeleton is not well understood. Here we identify the actin crosslinker Fimbrin as an essential force-sensitive regulator of cytoskeletal organization and cell adhesion in the *Drosophila* embryo. We show that Fimbrin is part of a mechanical feedback loop that amplifies mechanical forces at tricellular junctions and is required to recruit proteins that stabilize cell adhesion under tension (Fig. 5L). In the absence of Fimbrin, tricellular adhesion is disrupted under moderate levels of contractility and cells fail to reorganize actin and recruit junction-stabilizing proteins under force. Conversely, increased Fimbrin activity enhances the recruitment of myosin-II and junction-stabilizing proteins, indicating that Fimbrin promotes both the forces that exert stress on tricellular junctions and the junctional responses that resist this stress. This amplification step could allow mechanically regulated changes that are activated by different levels of tension to simultaneously reach the threshold for activation, enabling a coordinated, unified response to mechanical challenges to junctional stability.

The actin cytoskeleton undergoes several force-responsive changes *in vitro*, but how actin structures are influenced by physiological forces *in vivo* is not well understood. Our results demonstrate that Fimbrin-dependent actin crosslinking is an essential step in the activation of force responses during epithelial remodeling. In one model, Fimbrin could act as a mechanosensor, as other actin crosslinkers with Calponin-homology domains, such as α-actinin and filamin, can form force-sensitive catch bonds with actin *in vitro* (*33, 34*). Alternatively, actin filaments could act as mechanosensors in this process, undergoing conformational changes that recruit Fimbrin, as tensed actin filaments preferentially interact with other actin-binding proteins, including LIM domain proteins and α-catenin (*21, 27, 28*). In a third scenario, mechanical forces could promote a higher-order configuration of actin filaments that promotes Fimbrin binding, in a distinct, cytoskeletal network-level mechanism of force sensing. Indeed, the smaller size of Fimbrin, coupled with its ability to form compact actin networks with small interfilament spacing (*80-83*), may facilitate its ability to penetrate and reorganize the tightly packed, disordered actin networks characteristic of the actomyosin cortex, in contrast to larger actin crosslinkers that function in load-bearing structures such as stress fibers and muscle sarcomeres (*84, 85*). Once recruited, Fimbrin could generate mixed-polarity actin networks that trap myosin-II (*68, 80, 82, 83, 86*), align and strengthen actin networks to facilitate further force generation (*87, 88*), and drive the recruitment of myosin-II and other force-sensitive proteins (*29-32*). Mechanical feedback in actomyosin networks is critical for the spatiotemporally regulated forces that drive cell shape (*87, 89, 90*), epithelial remodeling (*70-76*), and tissue maintenance and repair under stress (*77, 78, 91*). Here we show that this feedback requires Fimbrin-dependent actin remodeling through a mechanism that acts directly on the actin cytoskeleton. As the cellular pathways that govern cell-surface mechanics converge on the actin cytoskeleton, tension-sensitive changes in actin architecture could provide a general strategy to integrate the pathways required to convert mechanical information into diverse cell behaviors *in vivo*.

## Acknowledgments

We thank Corey Elowsky and Hannah Ulman for the annotated images in Fig. 4E, Movie S4, and Movie S5, Michael Buszczak for the *pUASp-W-attB* vector, Ben Glick for monomeric superfolder Venus, Carolyn Ton for help with graphic illustration, and Rodrigo Fernandez-Gonzalez, Claire Looney, Lila Neahring, Nathan Shugarts Devanapally, Masako Tamada, Hannah Ulman, and Huapeng Yu for comments on the manuscript. Stocks generated by the Transgenic RNAi Project at Harvard Medical School and obtained from the Bloomington *Drosophila* Stock Center (NIH P4OODO18537) and the Kyoto Stock Center at the Kyoto Institute of Technology were used in this study. This work was funded by NIH/NIGMS R01 grant GM079340 to J.A.Z. and a Revson Foundation Senior Biomedical Fellowship to N.T. J.A.Z. is an investigator of the Howard Hughes Medical Institute.

## Notes

### Competing Interest Statement

The authors have declared no competing interest.

## References

1. C. P. Heisenberg, Y. Bellaiche, Forces in tissue morphogenesis and patterning. Cell 153, 948–962 (2013).

2. F. Martino, A. R. Perestrelo, V. Vinarsky, S. Pagliari, G. Forte, Cellular Mechanotransduction: From Tension to Function. Front Physiol 9, 824 (2018).

3. A. S. Yap, K. Duszyc, V. Viasnoff, Mechanosensing and Mechanotransduction at Cell-Cell Junctions. Cold Spring Harb Perspect Biol 10, a028761 (2018).

4. H. Wolfenson, B. Yang, M. P. Sheetz, Steps in Mechanotransduction Pathways that Control Cell Morphology. Annu Rev Physiol 81, 585–605 (2019).

5. T. Higashi, A. L. Miller, Tricellular junctions: how to build junctions at the TRICkiest points of epithelial cells. Mol Biol Cell 28, 2023–2034 (2017).

6. F. Bosveld, Z. Wang, Y. Bellaiche, Tricellular junctions: a hot corner of epithelial biology. Curr Opin Cell Biol 54, 80–88 (2018).

7. T. E. Vanderleest et al., Vertex sliding drives intercalation by radial coupling of adhesion and actomyosin networks during Drosophila germband extension. eLife 7, e34586 (2018).

8. K. E. Cavanaugh et al., Force-dependent intercellular adhesion strengthening underlies asymmetric adherens junction contraction. Curr Biol 32, 1986–2000 e1985 (2022).

9. L. van den Goor, J. Iseler, K. M. Koning, A. L. Miller, Mechanosensitive recruitment of Vinculin maintains junction integrity and barrier function at epithelial tricellular junctions. Current Biology 34, 4677-4691.e4675 (2024).

10. H. H. Yu, J. A. Zallen, Abl and Canoe/Afadin mediate mechanotransduction at tricellular junctions. Science 370, eaba5528 (2020).

11. K. Z. Perez-Vale et al., Multivalent interactions make adherens junction-cytoskeletal linkage robust during morphogenesis. J Cell Biol 220, e202104087 (2021).

12. L. Sheppard et al., The alpha-Catenin mechanosensing M region is required for cell adhesion during tissue morphogenesis. J Cell Biol 222, e202108091 (2023).

13. W. Razzell, M. E. Bustillo, J. A. Zallen, The force-sensitive protein Ajuba regulates cell adhesion during epithelial morphogenesis. J Cell Biol 217, 3715–3730 (2018).

14. S. Rakshit, Y. Zhang, K. Manibog, O. Shafraz, S. Sivasankar, Ideal, catch, and slip bonds in cadherin adhesion. Proc Natl Acad Sci U S A 109, 18815–18820 (2012).

15. K. Manibog, H. Li, S. Rakshit, S. Sivasankar, Resolving the molecular mechanism of cadherin catch bond formation. Nat Commun 5, 3941 (2014).

16. Q. le Duc et al., Vinculin potentiates E-cadherin mechanosensing and is recruited to actin-anchored sites within adherens junctions in a myosin II-dependent manner. J Cell Biol 189, 1107–1115 (2010).

17. S. Yonemura, Y. Wada, T. Watanabe, A. Nagafuchi, M. Shibata, alpha-Catenin as a tension transducer that induces adherens junction development. Nat Cell Biol 12, 533–542 (2010).

18. R. M. Mege, N. Ishiyama, Integration of Cadherin Adhesion and Cytoskeleton at Adherens Junctions. Cold Spring Harb Perspect Biol 9, a028738 (2017).

19. C. D. Buckley et al., Cell adhesion. The minimal cadherin-catenin complex binds to actin filaments under force. Science 346, 1254211 (2014).

20. D. L. Huang, N. A. Bax, C. D. Buckley, W. I. Weis, A. R. Dunn, Vinculin forms a directionally asymmetric catch bond with F-actin. Science 357, 703–706 (2017).

21. L. Mei et al., Molecular mechanism for direct actin force-sensing by alpha-catenin. Elife 9, e62514 (2020).

22. D. N. Clarke, A. C. Martin, Actin-based force generation and cell adhesion in tissue morphogenesis. Curr Biol 31, R667–R680 (2021).

23. G. Salbreux, G. Charras, E. Paluch, Actin cortex mechanics and cellular morphogenesis. Trends Cell Biol 22, 536–545 (2012).

24. T. D. Pollard, Actin and Actin-Binding Proteins. Cold Spring Harb Perspect Biol 8, a018226 (2016).

25. P. Chugh, E. K. Paluch, The actin cortex at a glance. J Cell Sci 131, jcs186254 (2018).

26. T. M. Svitkina, Actin Cell Cortex: Structure and Molecular Organization. Trends Cell Biol 30, 556–565 (2020).

27. X. Sun et al., Mechanosensing through Direct Binding of Tensed F-Actin by LIM Domains. Dev Cell 55, 468–482 e467 (2020).

28. J. D. Winkelman, C. A. Anderson, C. Suarez, D. R. Kovar, M. L. Gardel, Evolutionarily diverse LIM domain-containing proteins bind stressed actin filaments through a conserved mechanism. Proc Natl Acad Sci U S A 117, 25532–25542 (2020).

29. C. Veigel, J. E. Molloy, S. Schmitz, J. Kendrick-Jones, Load-dependent kinetics of force production by smooth muscle myosin measured with optical tweezers. Nat Cell Biol 5, 980–986 (2003).

30. M. Kovacs, K. Thirumurugan, P. J. Knight, J. R. Sellers, Load-dependent mechanism of nonmuscle myosin 2. Proc Natl Acad Sci U S A 104, 9994–9999 (2007).

31. M. J. Greenberg, H. Shuman, E. M. Ostap, Inherent force-dependent properties of beta-cardiac myosin contribute to the force-velocity relationship of cardiac muscle. Biophys J 107, L41–L44 (2014).

32. J. Sung et al., Harmonic force spectroscopy measures load-dependent kinetics of individual human beta-cardiac myosin molecules. Nat Commun 6, 7931 (2015).

33. L. Rognoni, J. Stigler, B. Pelz, J. Ylanne, M. Rief, Dynamic force sensing of filamin revealed in single-molecule experiments. Proc Natl Acad Sci U S A 109, 19679–19684 (2012).

34. Y. Mulla et al., Weak catch bonds make strong networks. Nat Mater 21, 1019–1023 (2022).

35. M. M. Kozlov, A. D. Bershadsky, Processive capping by formin suggests a force-driven mechanism of actin polymerization. J Cell Biol 167, 1011–1017 (2004).

36. S. Ebrahim, B. Kachar, Myosin transcellular networks regulate epithelial apical geometry. Cell Cycle 12, 2931–2932 (2013).

37. J. M. Leerberg et al., Tension-sensitive actin assembly supports contractility at the epithelial zonula adherens. Curr Biol 24, 1689–1699 (2014).

38. B. R. Acharya et al., A Mechanosensitive RhoA Pathway that Protects Epithelia against Acute Tensile Stress. Dev Cell 47, 439–452 e436 (2018).

39. L. M. Hoffman, C. C. Jensen, A. Chaturvedi, M. Yoshigi, M. C. Beckerle, Stretch-induced actin remodeling requires targeting of zyxin to stress fibers and recruitment of actin regulators. Mol Biol Cell 23, 1846–1859 (2012).

40. D. Y. Z. Phua, X. Sun, G. M. Alushin, Force-activated zyxin assemblies coordinate actin nucleation and crosslinking to orchestrate stress fiber repair. bioRxiv, 2024.2005.2017.594765 (2024).

41. R. E. Stephenson et al., Rho Flares Repair Local Tight Junction Leaks. Dev Cell 48, 445–459 e445 (2019).

42. X. Wu, J. A. Hammer, ZEISS Airyscan: Optimizing Usage for Fast, Gentle, Super-Resolution Imaging. Methods Mol Biol 2304, 111–130 (2021).

43. A. C. Pare, J. A. Zallen, Cellular, molecular, and biophysical control of epithelial cell intercalation. Curr Top Dev Biol 136, 167–193 (2020).

44. N. Efimova, T. M. Svitkina, Branched actin networks push against each other at adherens junctions to maintain cell-cell adhesion. J Cell Biol 217, 1827–1845 (2018).

45. K. E. Kasza, S. Supriyatno, J. A. Zallen, Cellular defects resulting from disease-related myosin II mutations in Drosophila. Proc Natl Acad Sci U S A 116, 22205–22211 (2019).

46. M. V. de Arruda, S. Watson, C. S. Lin, J. Leavitt, P. Matsudaira, Fimbrin is a homologue of the cytoplasmic phosphoprotein plastin and has domains homologous with calmodulin and actin gelation proteins. J Cell Biol 111, 1069–1079 (1990).

47. V. Delanote, J. Vandekerckhove, J. Gettemans, Plastins: versatile modulators of actin organization in (patho)physiological cellular processes. Acta Pharmacol Sin 26, 769–779 (2005).

48. C. M. Lye, H. W. Naylor, B. Sanson, Subcellular localisations of the CPTI collection of YFP-tagged proteins in Drosophila embryos. Development 141, 4006–4017 (2014).

49. A. E. Adams, D. Botstein, D. G. Drubin, Requirement of yeast fimbrin for actin organization and morphogenesis in vivo. Nature 354, 404–408 (1991).

50. R. Taylor et al., Absence of plastin 1 causes abnormal maintenance of hair cell stereocilia and a moderate form of hearing loss in mice. Hum Mol Genet 24, 37–49 (2015).

51. W. Y. Ding et al., Plastin increases cortical connectivity to facilitate robust polarization and timely cytokinesis. J Cell Biol 216, 1371–1386 (2017).

52. E. Dor-On et al., T-plastin is essential for basement membrane assembly and epidermal morphogenesis. Sci Signal 10, eaal3154 (2017).

53. D. Krueger, T. Quinkler, S. A. Mortensen, C. Sachse, S. De Renzis, Cross-linker-mediated regulation of actin network organization controls tissue morphogenesis. J Cell Biol 218, 2743–2761 (2019).

54. D. Garbett et al., T-Plastin reinforces membrane protrusions to bridge matrix gaps during cell migration. Nat Commun 11, 4818 (2020).

55. J. M. Hill, S. Cai, M. D. Carver, D. G. Drubin, A role for cross-linking proteins in actin filament network organization and force generation. Proc Natl Acad Sci U S A 121, e2407838121 (2024).

56. S. Narumiya, T. Ishizaki, M. Uehata, Use and properties of ROCK-specific inhibitor Y-27632. Methods Enzymol 325, 273–284 (2000).

57. H. Ishihara et al., Calyculin A and okadaic acid: inhibitors of protein phosphatase activity. Biochem Biophys Res Commun 159, 871–877 (1989).

58. J. C. Yu, R. Fernandez-Gonzalez, Local mechanical forces promote polarized junctional assembly and axis elongation in Drosophila. Elife 5, e10757 (2016).

59. C. Bertet, L. Sulak, T. Lecuit, Myosin-dependent junction remodelling controls planar cell intercalation and axis elongation. Nature 429, 667–671 (2004).

60. J. A. Zallen, E. Wieschaus, Patterned gene expression directs bipolar planar polarity in Drosophila. Dev Cell 6, 343–355 (2004).

61. M. S. Hutson et al., Forces for morphogenesis investigated with laser microsurgery and quantitative modeling. Science 300, 145–149 (2003).

62. R. Farhadifar, J. C. Roper, B. Aigouy, S. Eaton, F. Julicher, The influence of cell mechanics, cell-cell interactions, and proliferation on epithelial packing. Curr Biol 17, 2095–2104 (2007).

63. B. Boettner et al., The AF-6 homolog canoe acts as a Rap1 effector during dorsal closure of the Drosophila embryo. Genetics 165, 159–169 (2003).

64. T. T. Bonello, K. Z. Perez-Vale, K. D. Sumigray, M. Peifer, Rap1 acts via multiple mechanisms to position Canoe and adherens junctions and mediate apical-basal polarity establishment. Development 145, dev157941 (2018).

65. K. Z. Perez-Vale, K. D. Yow, N. J. Gurley, M. Greene, M. Peifer, Rap1 regulates apical contractility to allow embryonic morphogenesis without tissue disruption and acts in part via Canoe-independent mechanisms. Mol Biol Cell 34, ar7 (2023).

66. K. E. Rothenberg, Y. Chen, J. A. McDonald, R. Fernandez-Gonzalez, Rap1 coordinates cell-cell adhesion and cytoskeletal reorganization to drive collective cell migration in vivo. Curr Biol 33, 2587–2601 e2585 (2023).

67. S. M. Brower, J. E. Honts, A. E. Adams, Genetic analysis of the fimbrin-actin binding interaction in Saccharomyces cerevisiae. Genetics 140, 91–101 (1995).

68. L. Mei et al., Structural mechanism for bidirectional actin cross-linking by T-plastin. Proc Natl Acad Sci U S A 119, e2205370119 (2022).

69. D. L. Farrell, O. Weitz, M. O. Magnasco, J. A. Zallen, SEGGA: a toolset for rapid automated analysis of epithelial cell polarity and dynamics. Development 144, 1725–1734 (2017).

70. J. T. Blankenship, S. T. Backovic, J. S. Sanny, O. Weitz, J. A. Zallen, Multicellular rosette formation links planar cell polarity to tissue morphogenesis. Dev Cell 11, 459–470 (2006).

71. R. Fernandez-Gonzalez, S. Simoes, J. C. Roper, S. Eaton, J. A. Zallen, Myosin II dynamics are regulated by tension in intercalating cells. Dev Cell 17, 736–743 (2009).

72. P. A. Pouille, P. Ahmadi, A. C. Brunet, E. Farge, Mechanical signals trigger Myosin II redistribution and mesoderm invagination in Drosophila embryos. Sci Signal 2, ra16 (2009).

73. K. P. Landsberg et al., Increased cell bond tension governs cell sorting at the Drosophila anteroposterior compartment boundary. Curr Biol 19, 1950–1955 (2009).

74. B. Monier, A. Pelissier-Monier, A. H. Brand, B. Sanson, An actomyosin-based barrier inhibits cell mixing at compartmental boundaries in Drosophila embryos. Nat Cell Biol 12, 60–69 (2010).

75. J. C. Yu, N. Balaghi, G. Erdemci-Tandogan, V. Castle, R. Fernandez-Gonzalez, Myosin cables control the timing of tissue internalization in the Drosophila embryo. Cells Dev 168, 203721 (2021).

76. A. Munjal, J. M. Philippe, E. Munro, T. Lecuit, A self-organized biomechanical network drives shape changes during tissue morphogenesis. Nature 524, 351–355 (2015).

77. A. B. Kobb, T. Zulueta-Coarasa, R. Fernandez-Gonzalez, Tension regulates myosin dynamics during Drosophila embryonic wound repair. J Cell Sci 130, 689–696 (2017).

78. M. Duda et al., Polarization of Myosin II Refines Tissue Material Properties to Buffer Mechanical Stress. Dev Cell 48, 245–260 e247 (2019).

79. A. C. Pare et al., A positional Toll receptor code directs convergent extension in Drosophila. Nature 515, 523–527 (2014).

80. A. Bretscher, Fimbrin is a cytoskeletal protein that crosslinks F-actin in vitro. Proc Natl Acad Sci U S A 78, 6849–6853 (1981).

81. J. D. Winkelman et al., Fascin- and alpha-Actinin-Bundled Networks Contain Intrinsic Structural Features that Drive Protein Sorting. Curr Biol 26, 2697–2706 (2016).

82. K. L. Weirich, S. Stam, E. Munro, M. L. Gardel, Actin bundle architecture and mechanics regulate myosin II force generation. Biophys J 120, 1957–1970 (2021).

83. R. Sakamoto, M. P. Murrell, F-actin architecture determines the conversion of chemical energy into mechanical work. Nat Commun 15, 3444 (2024).

84. J. M. Lopez-Gay et al., Apical stress fibers enable a scaling between cell mechanical response and area in epithelial tissue. Science 370, eabb2169 (2020).

85. L. A. B. Fisher et al., Filamin protects myofibrils from contractile damage through changes in its mechanosensory region. PLoS Genet 20, e1011101 (2024).

86. C. L. Schwebach et al., Allosteric regulation controls actin-bundling properties of human plastins. Nat Struct Mol Biol 29, 519–528 (2022).

87. Y. Ren et al., Mechanosensing through cooperative interactions between myosin II and the actin crosslinker cortexillin I. Curr Biol 19, 1421–1428 (2009).

88. A. M. Lynch et al., TES-1/Tes and ZYX-1/Zyxin protect junctional actin networks under tension during epidermal morphogenesis in the C. elegans embryo. Curr Biol 32, 5189–5199 e5186 (2022).

89. J. C. Effler et al., Mitosis-specific mechanosensing and contractile-protein redistribution control cell shape. Curr Biol 16, 1962–1967 (2006).

90. N. Taneja et al., Precise Tuning of Cortical Contractility Regulates Cell Shape during Cytokinesis. Cell Rep 31, 107477 (2020).

91. R. Priya et al., Feedback regulation through myosin II confers robustness on RhoA signalling at E-cadherin junctions. Nat Cell Biol 17, 1282–1293 (2015).

