## Supplementary Material for "Actin crosslinking is required for force sensing at tricellular junctions"

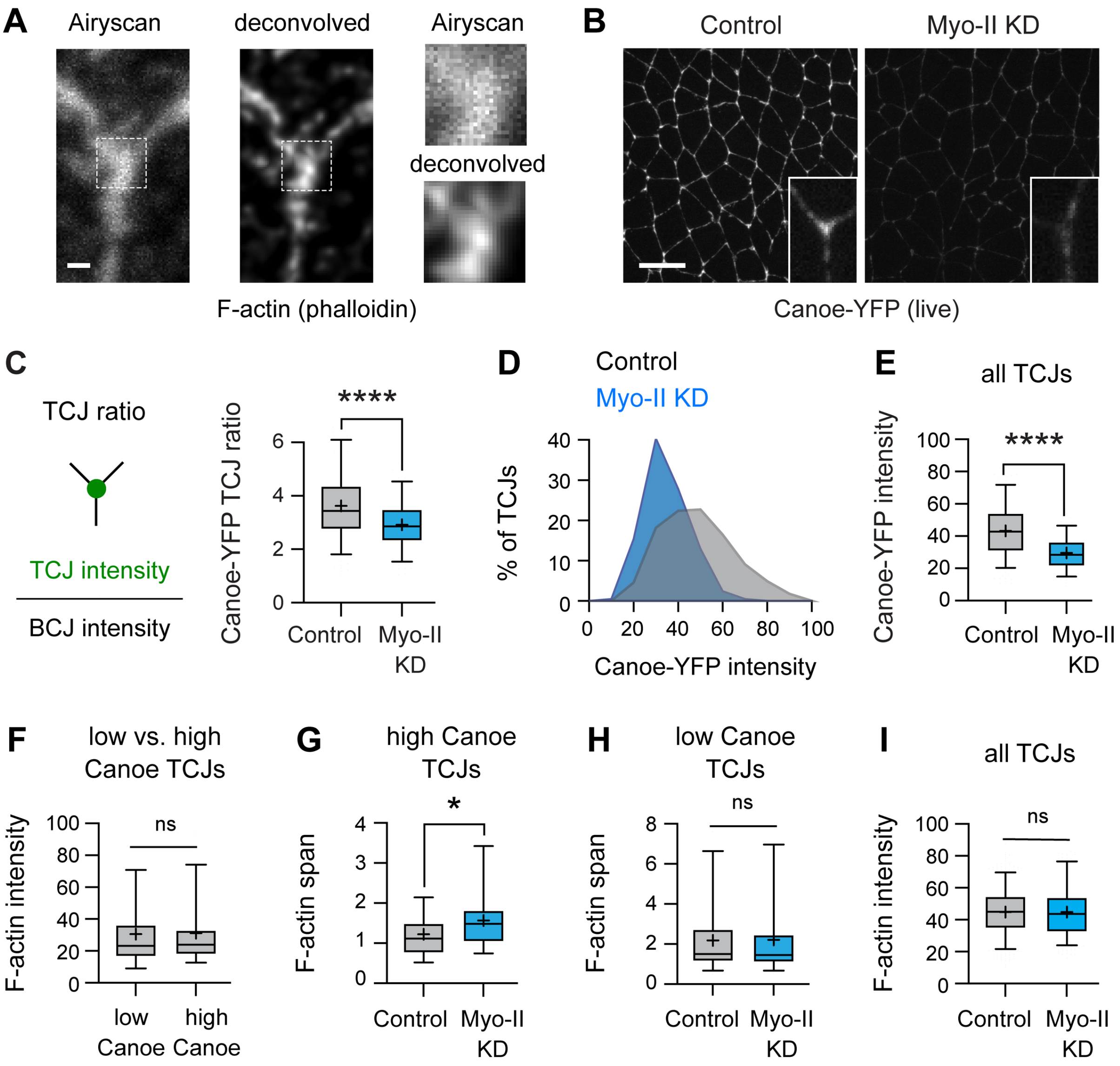


**Figure S1. Additional analysis of actin reorganization at tricellular junctions under tension**
**(A)** Increased resolution in Airyscan images after deconvolution. Localization of F-actin (phalloidin) at a single tricellular junction (TCJ). Boxes in left panels indicate regions shown on the right. **(B, C)** Localization (B) and TCJ ratio (TCJ intensity divided by the mean intensity of the three connected BCJs) (C) of Canoe-YFP in control (Gal4 only) and Myo-II KD embryos. **(D, E)** Canoe-YFP intensity at all TCJs in control and Myo-II KD embryos. **(F)** F-actin intensity at low and high Canoe TCJs in wild-type (WT) embryos (plotted for the TCJs analyzed in Fig. 1C). **(G, H)** F-actin span at high Canoe TCJs (top quartile, G) and low Canoe TCJs (bottom quartile, H) in control and Myo-II KD embryos (data from Figs. 1E and 1F). **(I)** F-actin intensity at all TCJs in control and Myo-II KD embryos. Live stage 7-8 embryos are shown in (B-E) and fixed stage 7-8 embryos are shown in (A) and (F-I), oriented ventral down in (B), 208-286 TCJs in 5-6 embryos/genotype in (C-H), 521-638 TCJs in 5-6 embryos/genotype in (I). Boxes, 25$^{th}$-75$^{th}$ percentile; whiskers, 5$^{th}$-95$^{th}$ percentile; horizontal line, median; +, mean. *, $p<0.05$, ***, $p<0.001$, ****, $p<0.0001$ (unpaired t-test with Welch's correction). Bars, 0.5 μm in (A), 10 μm in (B).

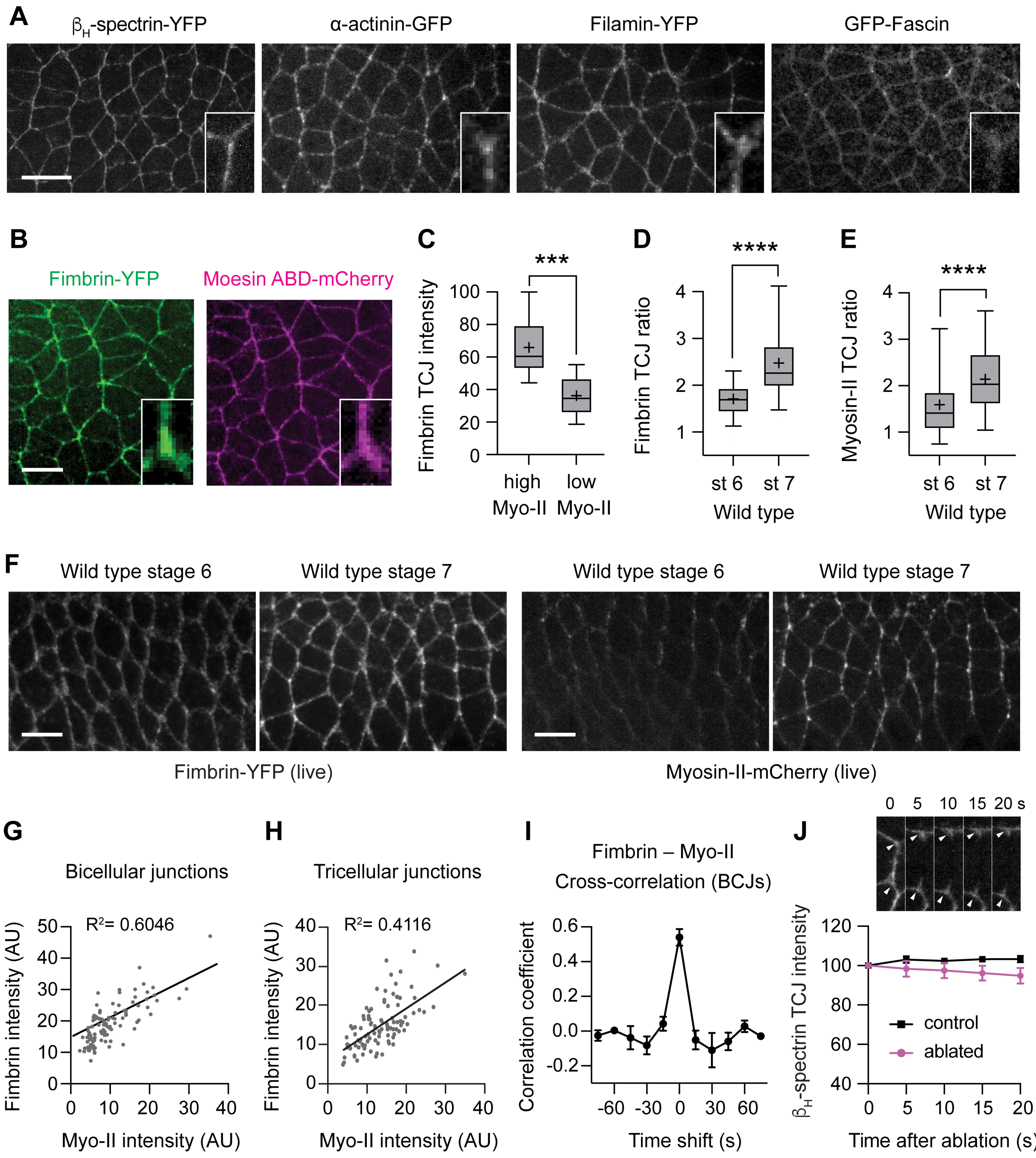


**Figure S2. A screen for actin crosslinkers that localize to tricellular junctions**

**(A, B)** Localization of βH-spectrin-YFP (karst), α-actinin-GFP, Filamin-YFP (cheerio), and GFP-Fascin (singed), Fimbrin-YFP, and F-actin (Moesin actin-binding domain-mCherry) in wild-type (WT) embryos. Insets, close-ups of single tricellular junctions (TCJs). **(C)** Fimbrin-YFP intensity at high myosin-II TCJs (top quartile) and low myosin-II TCJs (bottom quartile) in WT embryos. **(D-F)** Localization (F) and TCJ ratios (D,E) of Fimbrin-YFP and myosin-II-mCherry in WT embryos at indicated stages. **(G, H)** Spatial correlation between Fimbrin-YFP and myosin-II-mCherry intensity at bicellular junctions (BCJs) (G) and TCJs (H). $R^2$ values calculated on dataset pooled from multiple embryos. **(I)** Temporal correlation between Fimbrin-YFP and myosin-II-mCherry intensity at BCJs. Maximum correlation at a time shift of 0 s indicates no time delay between the recruitment of myosin-II-mCherry and Fimbrin-YFP. **(J)** $\beta_H$-spectrin-YFP localization at TCJs after edge ablation, normalized to 100% prior to ablation (t=0). Arrowheads, TCJs before (0 s) and after (5-20 s) edge ablation. p=0.058 (unpaired t-test with Welch's correction) at 20 s. Live stage 7-8 embryos are shown in (A), (B), (E), and (G-J) and live stage 6-7 embryos are shown in (C), (D), and (F), oriented ventral down in (A), (B), and (F), 100-106 TCJs in 3-5 embryos/condition in (C-E), 117 BCJs and 100 TCJs in 3-5 embryos in (G) and (H), 24 BCJs in 3 embryos in (I), 26-39 TCJs in 11 embryos in (J). Mean±SEM in (I) and (J). Boxes, $25^{th}$-$75^{th}$ percentile; whiskers, $5^{th}$-$95^{th}$ percentile; horizontal line, median; +, mean. ***, p<0.001, ****, p<0.0001 (unpaired t-test with Welch's correction). Bars, 10 $\mu$m.

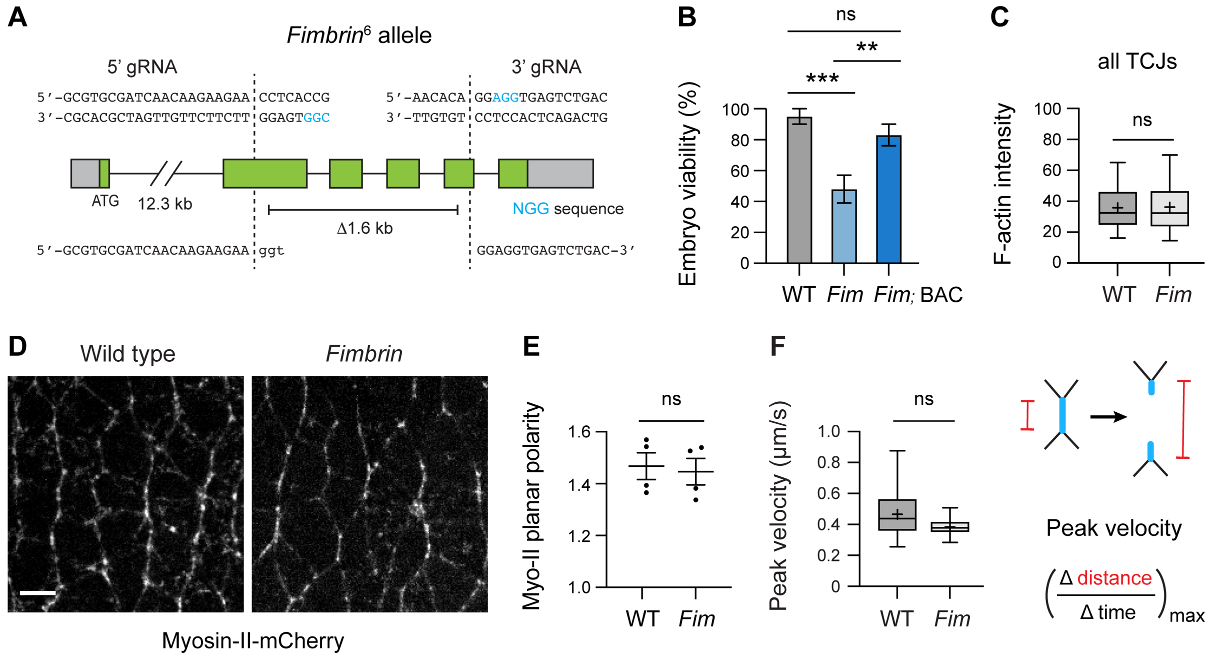


**Figure S3. Generation and analysis of *Fimbrin* mutant embryos**

**(A)** CRISPR-Cas9 editing of the *Fimbrin* locus. A 1.6 kb deletion was generated using gRNAs targeting exons 2 and 5. PAM sequences, blue. Bottom sequences, deletion breakpoints. **(B)** Embryo viability (the percentage of embryos that hatched into larvae) in wild type (WT), *Fimbrin* mutant (*Fim*) embryos, and *Fim* mutants with a bacterial artificial chromosome (BAC) containing WT *Fimbrin*. **(C)** F-actin intensity at all TCJs in WT and *Fim* embryos. **(D, E)** Myosin-II-GFP localization (D) and planar polarity (E) (ratio of the mean edge intensity parallel to the dorsal-ventral axis to the mean edge intensity perpendicular to this axis) in WT and *Fim* embryos. **(F)** Peak retraction velocity after ablation of isolated vertical edges (oriented within 30° of the dorsal-ventral axis) in WT and *Fim* embryos. Fixed stage 7 embryos are shown in (C), live stage 7 embryos are shown in (D) and (E), and live stage 7-8 embryos are shown in (F), oriented ventral down in (D), 60-117 embryos/genotype in (B), 216-233 TCJs in 5 embryos/genotype in (C), 530-641 edges in 4 embryos/genotype in (E), 7-14 edges/genotype in (F). Mean±SEM in (B) and (E). Boxes, 25$^{th}$-75$^{th}$ percentile; whiskers, 5$^{th}$-95$^{th}$ percentile; horizontal line, median; +, mean. ns, not significant, *, $p<0.05$, ***, $p<0.001$ (unpaired t-test with Welch's correction). Bars, 5 µm.

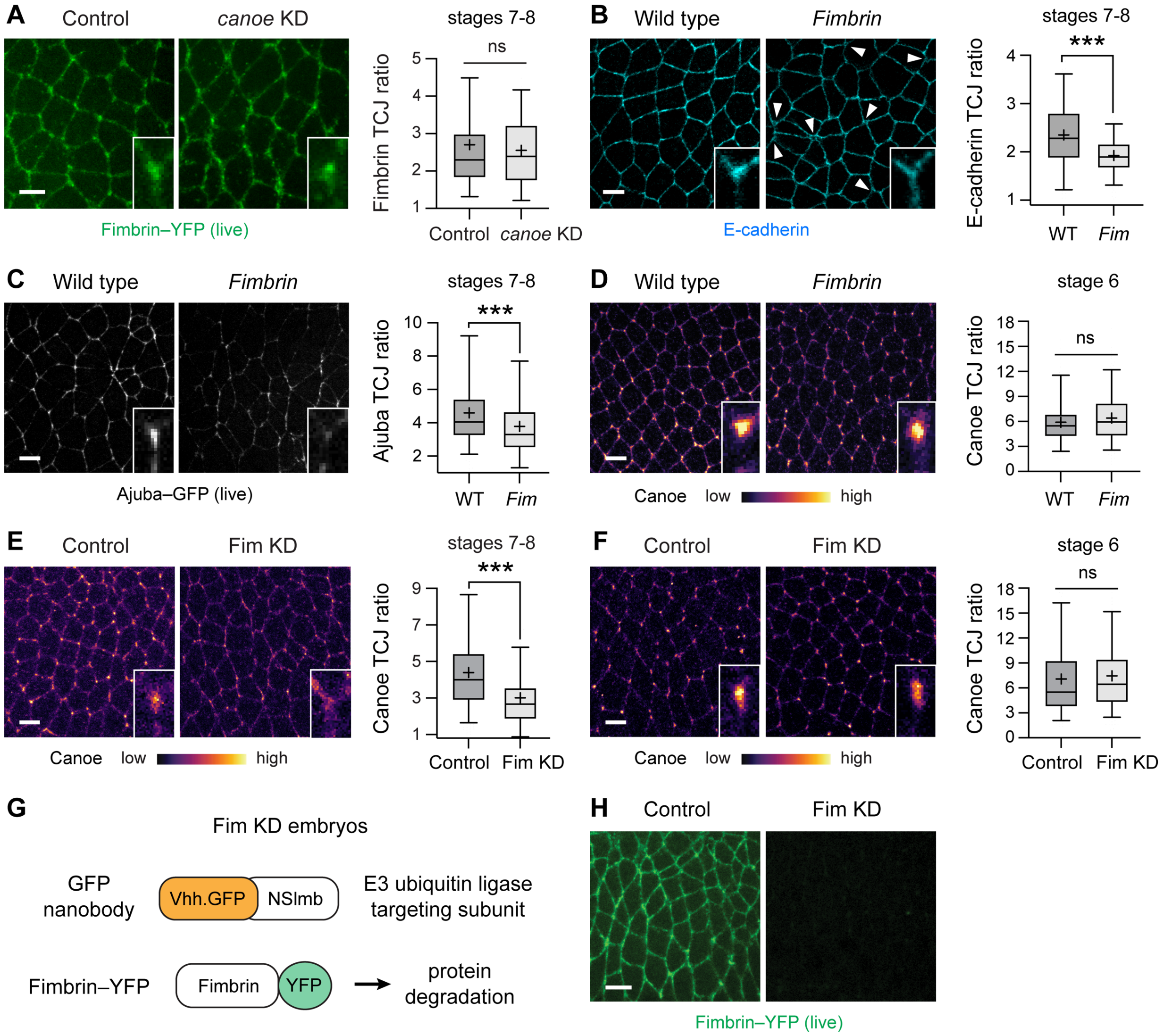


**Figure S4. Fimbrin is required to recruit multiple force-sensitive proteins to tricellular junctions** **(A-D)** Localization and TCJ ratios of Fimbrin-YFP (A), E-cadherin (B), Ajuba-GFP (C), and Canoe (D) in Control (Gal4 only), *canoe* KD, wild-type (WT), and *Fimbrin* mutant (*Fim*) embryos at stage 7 (A-C) or stage 6 (D). **(E, F)** Localization of Canoe in Control (Gal4 only) and Fim KD embryos at stage 7 (E) or stage 6 (F). **(G, H)** Proteasomal degradation of Fimbrin-YFP by expressing the E3 ubiquitin ligase targeting subunit fused to a nanobody targeting GFP (G) efficiently depletes Fimbrin-YFP in stage 7 Fim KD embryos (H). Embryos with >70% reduction of Fimbrin-YFP intensity compared with control embryos were included in the analysis. Live stage 7-8 embryos are shown in (A), (C) and (H), fixed stage 7-8 embryos are shown in (B) and (E), fixed stage 6 embryos are shown in (D) and (F), oriented ventral down, 179-378 TCJs in 3-11 embryos/genotype. Boxes, 25$^{th}$-75$^{th}$ percentile; whiskers, 5$^{th}$-95$^{th}$ percentile; horizontal line, median; +, mean. ns not significant, ***, p<0.0001 (unpaired t-test with Welch's correction). Bars, 5 μm.

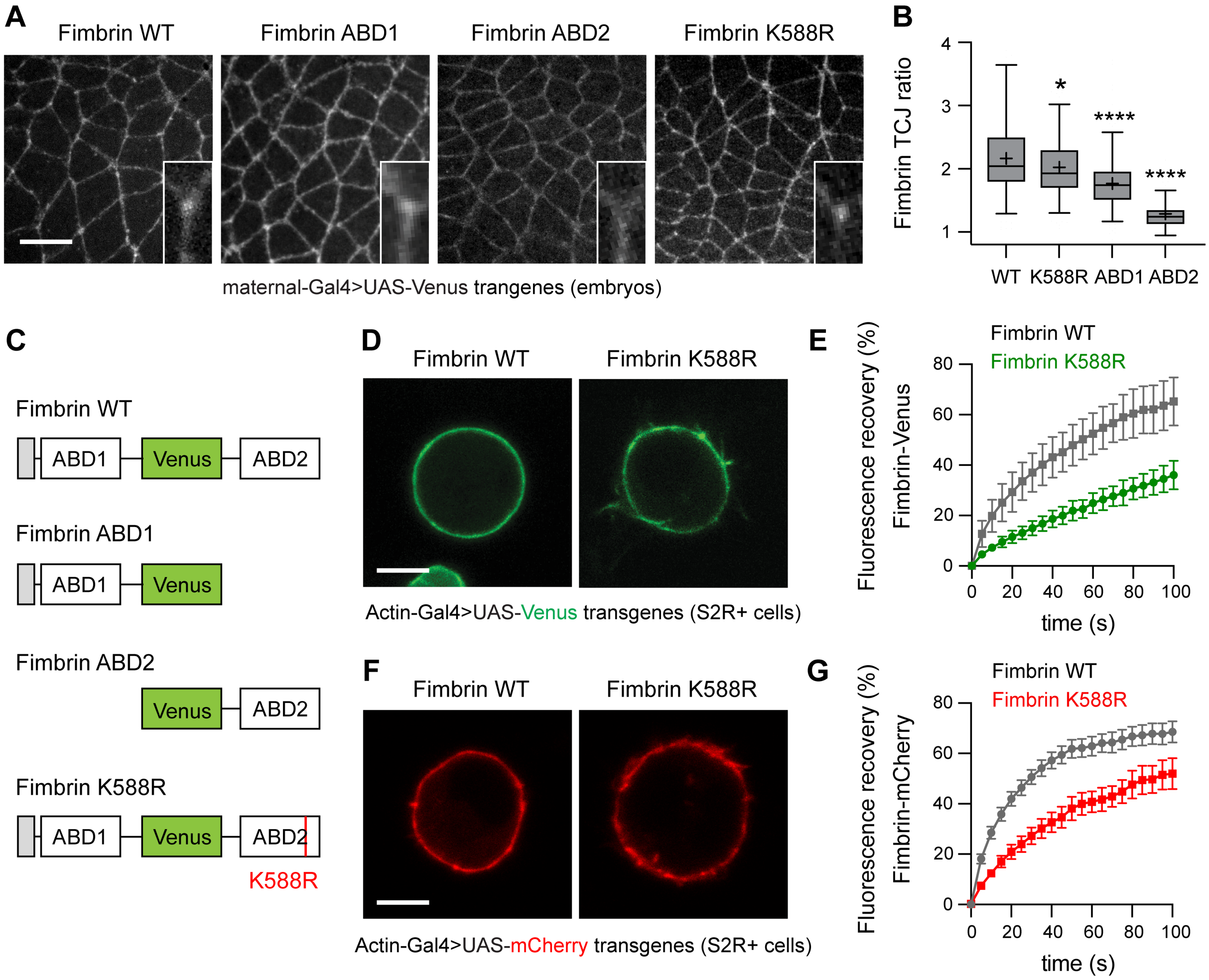


**Figure S5. Identification of sequences that modulate Fimbrin localization and dynamics**
**(A, B)** Localization (A) and TCJ ratios (B) of Venus-tagged Fimbrin transgenes. **(C)** Venus-tagged Fimbrin variants. Gray box, EF hand domain. ABD, actin-binding domain. **(D-G)** Localization (D, F) and fluorescence recovery after photobleaching (E, G) of Venus-tagged wild-type Fimbrin and Fimbrin$^{K588R}$ (D and E) and mCherry-tagged wild-type Fimbrin and Fimbrin$^{K588R}$ (F and G). $p<0.0001$ at 100 s in (E), $p<0.05$ at 100 s in (G) (unpaired t-test with Welch's correction). Live stage 7-8 embryos are shown in (A) and (B), oriented ventral down in (A), live S2R+ cells are shown in (D-G), 169-218 TCJs in 4-6 embryos/genotype in (B), 6-7 S2R+ cells/condition in (E) and (G). Mean±SEM in (E) and (G). Boxes, 25$^{th}$-75$^{th}$ percentile; whiskers, 5$^{th}$-95$^{th}$ percentile; horizontal line, median; +, mean. *, $p<0.05$, ****, $p<0.0001$ (unpaired t-test with Welch's correction). Bars, 10 μm in (A), 5 μm in (D) and (F).

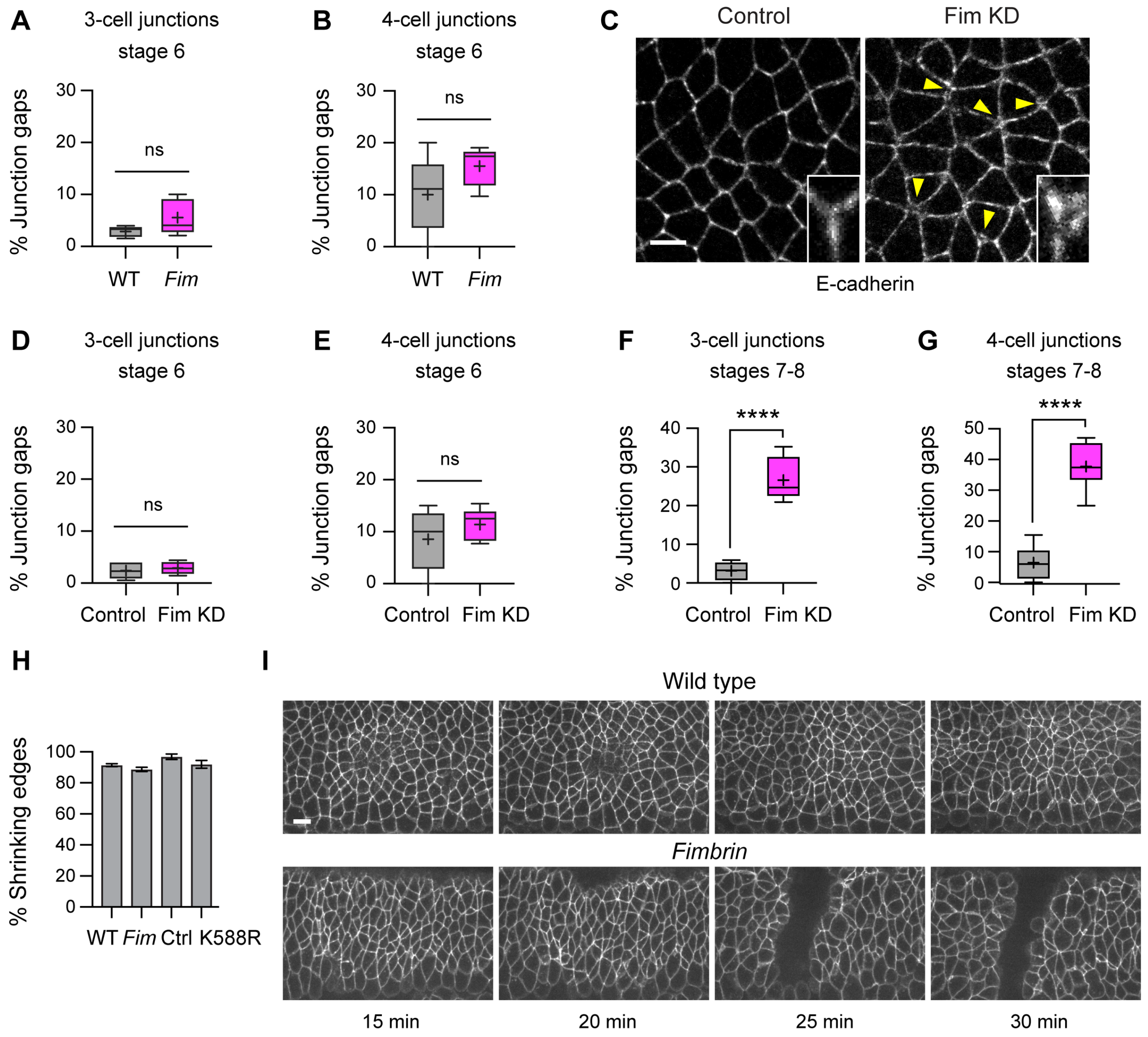


**Figure S6. Effects of Fimbrin on cell adhesion and behavior**
**(A, B)** Percentage of tricellular junctions (TCJs) (A) and 4-cell junctions (B) with gaps in wild-type (WT) and *Fimbrin* mutant (*Fim*) embryos at stage 6. **(C)** E-cadherin localization in control (Gal4 only) and Fim KD embryos. Arrowheads, gaps in E-cadherin signal. Insets, close-ups of single TCJs. **(D-G)** Percentage of TCJs (D and F) and 4-cell junctions (E and G) with gaps in control and Fim KD embryos at stage 6 (D and E) and stage 7 (F and G). **(H)** Percentage of vertical edges that shrink to a vertex in WT, *Fim*, control, and Fimbrin$^{K588R}$ embryos. **(I)** Aberrant grooves were observed in *Fim* embryos. Fixed stage 6 embryos are shown in (A), (B), (D), and (E), fixed stage 7-8 embryos are shown in (C), (F) and (G), live stage 7-8 embryos are shown in (H) and (I), oriented ventral down in (C) and (I), 588-1,002 TCJs and 83-138 4-cell junctions in 5-8 embryos/genotype in (A), (B) and (D-G), 93-127 edges in 3-4 embryos/genotype in (H). Boxes, 25$^{th}$-75$^{th}$ percentile; whiskers, 5$^{th}$-95$^{th}$ percentile; horizontal line, median; +, mean. ns, not significant, ****, $p<0.0001$ (unpaired t-test with Welch's correction). Bars, 5 $\mu$m in (C), 10 $\mu$m in (I).

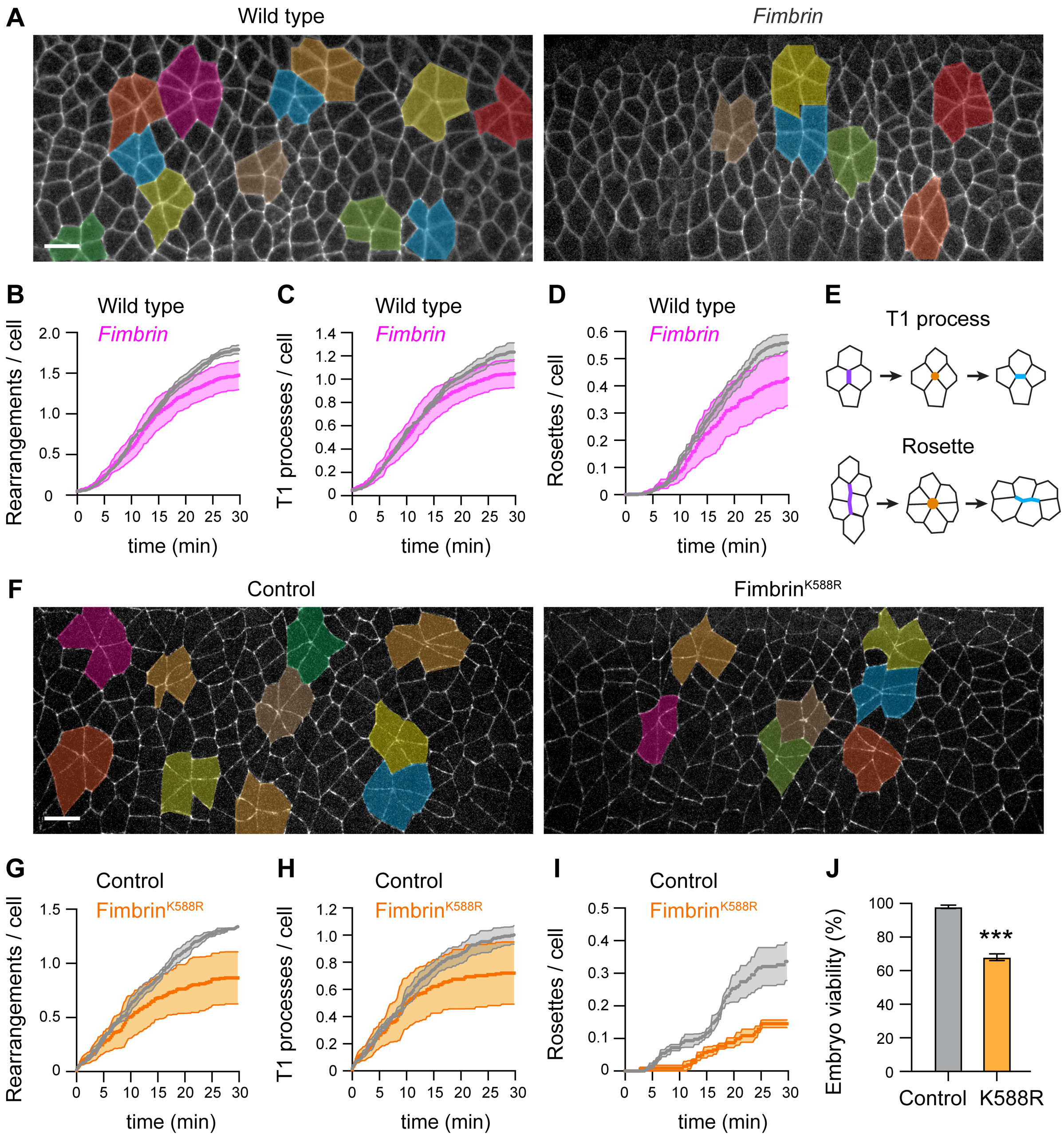


**Figure S7. Fimbrin regulates cell rearrangement during axis elongation**
**(A)** Stills from movies of wild-type (WT) and *Fimbrin* mutant (*Fim*) embryos at 17.5 min after the onset of elongation in stage 7. Cells in rosettes are highlighted. **(B-D)** Cumulative total cell rearrangements (B), T1 processes (C), and rosettes (D) per cell in WT and *Fim* embryos. **(E)** Cell rearrangement schematics. Edges oriented parallel to the dorsal-ventral axis (vertical edges, purple) contract to form a 4-cell junction or rosette (orange) that resolves through the formation of edges perpendicular to this axis (blue). **(F)** Stills from movies of control (Gal4 only) and Fimbrin$^{K588R}$ embryos at 17.5 min after the onset of elongation. Cells in rosettes are highlighted. **(G-I)** Cumulative total cell rearrangements (G), T1 processes (H), and rosettes (I) per cell in control and Fimbrin$^{K588R}$ embryos. **(J)** Embryo viability (the percentage of embryos that hatched into larvae) in control and Fimbrin$^{K588R}$ embryos. Live stage 7-8 embryos are shown in (A-I), oriented ventral down, 89-285 cells in 3-4 embryos/genotype in (B-D) and (G-I), 124-150 embryos in 3 replicates/genotype in (J). Mean±SEM in all panels. Bars, 10 μm.

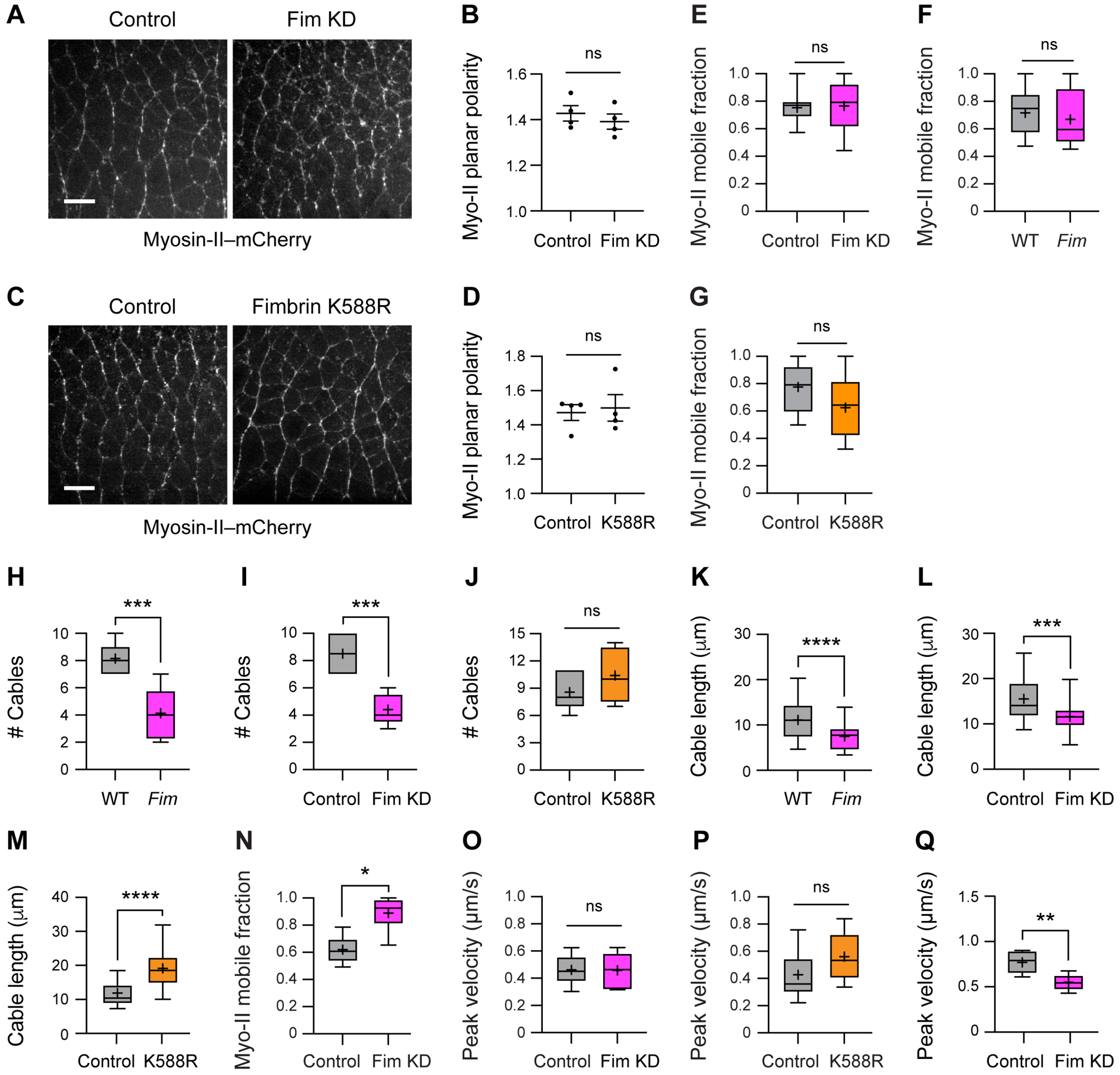


**Figure S8. Effects of Fimbrin on myosin-II localization, organization and activity**
**(A-D)** Myosin-II-mCherry localization (A and C), and planar polarity (B and D) in control (Gal4 only), Fim KD, and Fimbrin$^{K588R}$ embryos. Fimbrin$^{K588R}$, larger view of cropped version shown in Fig. 5G. **(E)** Myosin-II-mCherry mobile fraction (fraction of recovered myosin-II signal 35 s after photobleaching) at isolated vertical edges (oriented within 30° of the dorsoventral axis) in control and Fim KD embryos. (**F, G)** Myosin-II-GFP mobile fraction at isolated vertical edges in wild-type (WT) and *Fimbrin* mutant (*Fim*) (F) and control and Fimbrin$^{K588R}$ (G) embryos. **(H-M)** Myosin-II cable number (H-J) and length (K-M) in the indicated genotypes. **(N)** Myosin-II-mCherry mobile fraction in cables in control and Fim KD embryos. **(O, P)** Peak retraction velocity after ablation of isolated vertical edges in control, Fim KD, and Fimbrin$^{K588R}$ embryos. **(Q)** Peak retraction velocity after ablation of myosin-II cables in control and Fim KD embryos. Live stage 7-8 embryos are shown in all panels, oriented ventral down in (A) and (C), 639-884 edges in 4 embryos/genotype in (B) and (D), 6-15 cables or edges/genotype in (E-G) and (N-Q), 22-60 cables in 5-9 embryos/genotype in (H-M). Mean±SEM in (B) and (F). Boxes, 25$^{th}$-75$^{th}$ percentile; whiskers, 5$^{th}$-95$^{th}$ percentile; horizontal line, median; +, mean. ns, not significant, *, $p<0.05$, **, $p<0.01$, ***, $p<0.001$, ****, $p<0.0001$ (unpaired t-test with Welch's correction). Bars, 10 μm.

**Supplementary Movie Legends**

**Movie S1**. **Fimbrin-YFP recruitment to tricellular junctions in wild type.** Time-lapse movie of Fimbrin-YFP (green) and E-cadherin-mCherry (magenta) during the contraction phase after apical ablation. t=0 s, time of maximal area expansion. Dimensions, 40 μm x 20 μm.

**Movie S2. Canoe-YFP recruitment to tricellular junctions in wild type.** Time-lapse movie of Canoe-YFP (mpl-inferno look-up table) during the contraction phase after apical ablation. t=0 s, time of maximal area expansion. Dimensions, 20 μm x 20 μm.

**Movie S3. Canoe-YFP recruitment to tricellular junctions in *Fimbrin.*** Time-lapse movie of Canoe-YFP (mpl-inferno look-up table) during the contraction phase after apical ablation. t=0 s, time of maximal area expansion. Dimensions, 20 μm x 20 μm.

**Movie S4. Cell rearrangement rate in wild type and *Fimbrin*.** Time-lapse movie of wild-type (top) and *Fimbrin* (bottom) embryos expressing Spider-GFP during axis elongation. T1 processes are color-coded by vertex duration. Anterior left, ventral down. Dimensions/embryo, 110 μm x 65 μm.

**Movie S5. Cell rearrangement rate in control and Fimbrin$^{K588R}$.** Time-lapse movie of control (top) and Fimbrin-K588R (bottom) embryos expressing β-catenin-GFP during axis elongation. T1 processes are color-coded by vertex duration. Anterior left, ventral down. Dimensions/embryo, 104 μm x 56 μm.

**Movie S6. Myosin-II-mCherry recruitment to tricellular junctions in wild type.** Time-lapse movie of myosin-II-mCherry (cyan) and Spider-GFP (magenta) during the contraction phase after apical ablation. t=0 s, time of maximal area expansion. Dimensions, 40 μm x 20 μm.

**Movie S7. Myosin-II-mCherry recruitment to tricellular junctions in *Fimbrin*.** Time-lapse movie of myosin-II-mCherry (cyan) and Spider-GFP (magenta) during the contraction phase after apical ablation. t=0 s, time of maximal area expansion. Dimensions, 40 μm x 20 μm.

### Materials and Methods

### Antibodies and Reagents

| Antibody/Reagent | Manufacturer/Source | Concentration | Identifiers |
|---|---|---|---|
| mouse anti-$\alpha$-spectrin (3A9) | Developmental Studies Hybridoma Bank | 1:50 | Antibody registry ID: AB_528473 |
| rabbit anti-Canoe | (*1*) | 1:250 | N/A |
| rat anti-E-cadherin | (*2*), Developmental Studies Hybridoma Bank (DCAD2) | 1:50 (concentrated) | Antibody registry ID: AB_528120 |
| rabbit anti-GFP | Torrey Pines | 1:150 | NC9589665; RRID: AB_10013661 |
| GFP nanobooster-488 | Chromotek | 1:100 | gb2AF488; RRID: AB_2827573 |
| AlexaFluor-488 phalloidin | Thermo Fisher | 1:200 | A12379 |
| AlexaFluor-568 phalloidin | Thermo Fisher | 1:200 | A12380 |
| AlexaFluor-647 phalloidin | Thermo Fisher | 1:200 | A22287 |

### Fly stocks

| Stock | Reference |
|---|---|
| *$\alpha$-actinin–GFP* | (*3*) |
| *Ajuba–GFP BAC (III)* | (*4*) |
| *$\beta$-catenin–GFP (arm-arm–GFP) (II)* | (*5*), Bloomington *Drosophila* Stock Center (BDSC) 8556 |
| *$\beta$-catenin–GFP (arm-arm–GFP) (III)* | (*5*), BDSC 8555 |
| *$\beta_H$-spectrin–YFP* (*kst*$^{CPTI002266}$) | (*6*), Kyoto *Drosophila* Stock Center 115285 |
| *Canoe–YFP* (*cno*$^{CPTI000590}$) | (*6*), Kyoto *Drosophila* Stock Center 115111 |
| *E-cadherin–mCherry (shg-shg-mCherry) (II)* | (*7*) |
| *Filamin–YFP* (*cheerio*$^{CPTI001399}$) | (*6*), Kyoto *Drosophila* Stock Center 115514 |
| *Fimbrin–YFP (fim$^{CPTI100066}$)* | (*6*), Kyoto *Drosophila* Stock Center 115092 |
| *mat$\alpha$tub67 Gal4 (II)* | gift of Daniel St Johnston |
| *mat$\alpha$tub15 Gal4 (III)* | gift of Daniel St Johnston |
| *myosin-II–GFP (sqh-sqh–GFP) (II)* | (*8*) |
| *myosin-II–GFP (sqh-sqh–GFP) (III)* | (*8*) |
| *myosin-II–mCherry (sqh-sqh–mCherry) (II)* | (*9*) |
| *myosin-II–mCherry (sqh-sqh–mCherry) (III)* | (*9*), BDSC 59024 |
| *nos-Cas9 (III)* | (*10*), BDSC 78782 |
| *Spider–GFP* | gift of Alain Debec (Institut Jacques Monod) |
| *sqh-gap43–mCherry (III)* | (*11*) |

| | |
|---|---|
| *sqh-MoesinABD–mCherry (II)* | (*12*), BDSC 35520 |
| *sqh-Utrophin–mCherry (II)* | (*13*) |
| *UAS-canoe RNAi GL00633 (II)* | (*14*), BDSC 38194 |
| *UAS-GFP–Fascin* | (*15*) |
| *UASp-Nslmb-VHH–GFP* | (*16*), BDSC 38422 |
| *UAS-zipper RNAi GL00623 (II)* | (*14*), BDSC 37480 |
| *Fimbrin*$^{6}$*, FRT19A/FM7, sn* | This study |
| *UASp-Fim-Venus (attP2)* (internal tag) | This study |
| *UASp-Fim-mCherry (attP2)* (internal tag) | This study |
| *UASp-Fim*$^{K588R}$*-Venus (attP2)* (internal tag) | This study |
| *UASp-Fim*$^{K588R}$*-mCherry (attP2)* (internal tag) | This study |
| *UASp-Fim-ABD1-Venus (attP2)* | This study |
| *UASp-Venus-Fim-ABD2 (attP2)* | This study |
| *U6-Fim-gRNA* (attP2) | This study |

**Recombinant DNA**

| | |
|---|---|
| *pUASp-Fim-Venus (internal tag)* | This study |
| *pUASp-Fim-mCherry (internal tag)* | This study |
| *pUASp-Fim*$^{K588R}$*-Venus (internal tag)* | This study |
| *pUASp-Fim*$^{K588R}$*-mCherry (internal tag)* | This study |
| *pUASp-Fim-ABD1-Venus* | This study |
| *pUASp-Venus-Fim-ABD2* | This study |
| *pCFD4-U6-Fim-gRNA* | This study |

**Primers and synthetic gene fragments**

| **Primer** | **Sequence** | **Purpose** |
|---|---|---|
| pCFD4 Fim gRNA #1 fwd | 5′-TATATAGGAAAGATATCCGGGTGAACTTCGTTCTCAAACTCCCGGTAGAGTTTTAGAGCTAGAAATAGCAAG-3′ | 5' cut site for the *Fimbrin* null allele |
| pCFD4 Fim gRNA #2 rev | 5′-ATTTTAACTTGCTATTTCTAGCTCTAAAACGTGTTCCGCTAGTGCGCACCGACGTTAAATTGAAAATAGGTC-3′ | 3' cut site for the *Fimbrin* null allele |
| Fim-RA-ABD1-fwd | 5′-AAAGCAGGCTCCGCGGCCGCCCCCTTCACCATGGCAACACTTAACAAATTCACAAAAACG-3′ | to amplify *Fimbrin* codons 1-395 for *pUASp-Fim-Venus* and *pUASp-Fim-ABD1-Venus* |
| ABD1-3' msVenus-5'-rev | 5′-GCCCATTGAGTTCATCCAATTGCGGTAGGTCCCGCTGGAGCCGGAGC-3′ | to amplify *Fimbrin* codons 1-395 for *pUASp-Fim-Venus* |
| Fim ABD2 5' fwd | 5′-ACCTACCGCAATTGGATGAACTCAA-3′ | to amplify *Fimbrin* codons 396-640 for *pUASp-Fim-Venus* and *pUASp-Fim-mCherry* |
| Fim ABD2 3' rev | 5′-CAAGAAAGCTGGGTCGGCGCGCCCACCCTTCTAATTGCTGTTGTTGGCGGAGC-3′ | to amplify *Fimbrin* codons 396-640 for pUASp-Fim-Venus |

| | | |
|---|---|---|
| | | and *pUASp-Venus-Fim-ABD2*, and *pUASp-Fim-mCherry* |
| msVenus-5' fwd | ATGGATAGCACTGAGAGCCTGTTCACCG | to amplify Venus for *pUASp-Fim-Venus* |
| msVenus-3' ABD2-5' rev | 5′-GCCCATTGAGTTCATCCAATTGCGGTAGGTCCCGCTGGAGCCGGAGC-3′ | to amplify Venus for *pUASp-Fim-Venus* |
| ABD1-3' mCherry-5' rev | 5′-CATGTTATCCTCCTCGCCCTTGCTCACCATCTTCTCTTCGCGCGTCTCCT-3′ | to amplify *Fimbrin* codons 1-395 for *pUASp-Fim-mCherry* |
| mCherry-5' fwd | 5′- ATGGTGAGCAAGGGCGAGGAGGATAACATG-3′ | to amplify mCherry for *pUASp-Fim-mCherry* |
| mCherry-3' rev | 5′-GCCCATTGAGTTCATCCAATTGCGGTAGGTCTTGTACAGCTCGTCCATGCCG-3′ | to amplify mCherry for *pUASp-Fim-mCherry* |
| K588R gBlock rev | 5′-TAAGCTGCCAGATGAGAGCCAACGT-3′ | to amplify *Fimbrin* nt 1-1483 for *pUASp-Fim$^{K588R}$-Venus* and *pUASp-Fim$^{K588R}$-mCherry* |
| msVenus-5' UASp-5' fwd | 5′-AAAGCAGGCTCCGCGGCCGCCCCCTTCACCATGGATAGCACTGAGAGCCTGTTCACC-3′ | to amplify Venus-ABD2 for *pUASp-Venus-Fim-ABD2* |
| mVenus-3' UASp-3' rev | CAAGAAAGCTGGGTCGGCGCGCCCACCCTTCTACCCGCTGGAGCCGG | to amplify ABD1-Venus for *pUASp-Fim-ABD1-Venus* |
| Fimbrin K588R gene block | 5′-CGCTGACGTTGGCTCTCATCTGGCAGCTTATGCGTGCCTACACCCTGTCCATTCTGTCCCGCTTGGCCAACACTGGCAACCCCATTATCGAGAAGGAGATCGTCCAGTGGGTGAATAACCGACTGTCGGAGGCAGGCAAACAGTCGCAGCTGCGTAACTTCAACGATCCGGCCATCGCCGATGGCAAGATCGTGATCGATCTGATCGATGCCATCAAGGAGGGCAGCATTAACTACGAGTTGGTGCGCACTAGCGGAACACAGGAGGATAACCTGGCCAATGCCAAGTATGCCATCTCCATGGCCCGCCGCATCGGCGCCCGTGTCTACGCCCTGCCCGAGGACATCACCGAGGTGAAGCCGAAAATGGTGATGACCGTTTTCGCCTGCATGATGGCCCTCGACTACGTGCCCAACATGGACAGTGTGGACCAGAACAACCACAACAGCTCCGCCAACAACAGCAATTAGAAGGGTGGGCGCGCCGACCCAGCTTTCTTG-3′ | synthetic gene fragment containing *Fimbrin* nt 1453-1923 and a 3' 30 bp overlap region with *pUASp-w-attB* |

**Plasmid construction**
All Venus-tagged constructs were constructed using monomeric superfolder Venus, designated as Venus throughout. To generate internally tagged full-length *pUASp-Fimbrin-Venus* and *pUASp-Fimbrin-mCherry* transgenes that recapitulate the internal site of the tag in the

endogenously tagged Fimbrin-YFP fly line (after amino acid K935 between the two actin-binding domains, mapped by (*17*)), the 5’ sequence encoding Fimbrin actin-binding domain 1 (ABD1) (aa 1-395) and the 3’ sequence encoding Fimbrin actin-binding domain 2 (ABD2) (aa 396-640) were PCR amplified from Fimbrin-RA clone LD05347 (*Drosophila* Genome Resource Center). Vector backbones were generated by linearizing the *pUASp-W-attB* vector with BamHI and MluI to excise the CAT/CmR-ccdB lethal gene cassette. The 5’ and 3’ *Fimbrin* PCR products, PCR-amplified *Venus* or *mCherry* fragments, and the linearized *pUASp-W-attB* vector were gel-purified using the GFX PCR DNA and Gel Band Purification kit (Cytiva) and assembled in a 4-part Gibson assembly reaction using NEBuilder HiFi DNA Assembly Master Mix (New England Biolabs, NEB). To generate *pUASp-FimABD1-Venus* and *pUASp-Venus-FimABD2*, fragments encoding ABD1-mVenus (Fimbrin aa 1-395) or Venus-ABD2 (Fimbrin aa 396-640) were PCR-amplified from the full-length internally tagged *pUASp-Fimbrin-Venus* plasmid and each separately combined with the linearized *pUASp-W-attB* vector in a 2-part Gibson assembly reaction.

To generate *pUASp-Fimbrin*$^{K588R}$*-mCherry* and *pUASp-Fimbrin*$^{K588R}$*-Venus*, a 3’ gene block fragment (*Fimbrin* nt 1,453-1,923 bearing the K588R AAG to CGC mutation) was synthesized (Integrated DNA Technologies). The 5’ region of *Fimbrin* (nt 1-1,483) with *Venus* or *mCherry* inserted before nt 1,136 was PCR-amplified from the *pUASp-Fimbrin-Venus* or *pUASp-Fimbrin-mCherry* vectors and each separately combined with the gene block and the linearized *pUASp-W-attB* vector in a 3-part Gibson assembly reaction.

The *pCFD4-U6-Fim-gRNA* plasmid was generated using previously described methods (*18*). Forward and reverse primers containing the two gRNA sequences were used to amplify the insert fragment from the undigested *pCD4-U6:gRNA-gRNA:U6* plasmid as a template. The *pCD4-U6:gRNA-gRNA:U6* plasmid was digested with BbsI to generate the vector backbone. The PCR product and linearized vector were gel-purified and combined in a 2-part Gibson assembly reaction.

### Fly crosses and genetics

The following fly stocks were obtained from the Bloomington *Drosophila* Stock Center (BDSC): *arm-arm-GFP* (II) and *arm-arm-GFP* (III) (β-catenin-GFP) (*5*), *nos-Cas9* (attP2), *sqh-MoesinABD-mCherry* (*12*), *UAS-canoe-shRNA* (GL00633) (attP40), *UASp-Nslmb-VHH-GFP* (*16*), and *UAS-zipper-shRNA* (GL00623) (attP40) (myosin-II shRNA). The following fly stocks were obtained from the Kyoto *Drosophila* Stock Center: $\beta_H$*-spectrin-YFP* (*kst*$^{CPTI002266}$) (*6*), *Canoe-YFP* (*cno*$^{CPTI000590}$) (*6*), *Filamin-YFP* (*cheerio*$^{CPTI001399}$) (*6*), and *Fimbrin-YFP* (*fim*$^{CPTI100066}$) (*6*). Other stocks used in this study were $\alpha$*-actinin-GFP* (*3*), *Ajuba-GFP* (*4*), *mat*$\alpha$*-tubulin67 (II)* and *mat*$\alpha$*-tubulin15 (III)* (gifts of D. St Johnson), *mat*$\alpha$*-tubulin15, sqh-gap43-mCherry/+* recombinant (*19*), *shg-shg-mCherry* (*7*), *Spider-GFP* (gift of Alain Debec), *sqh-sqh-GFP* (II) and *sqh-sqh-GFP* (III) (myosin-II-GFP) (*8*), *sqh-sqh-mCherry* (II) and *sqh-sqh-mCherry* (III) (myosin-II-mCherry) (*9*), *sqh-Utrophin-mCherry* (*13*), and *UAS-GFP-Fascin* (*15*). The following stocks were generated in this study: *Fimbrin*$^{6}$, *UASp-Fim-Venus, UASp-Fim-mCherry*, *UASp-Fim*$^{K588R}$*-Venus*, *UASp-Fim*$^{K588R}$*-mCherry, UASp-Fim-ABD1-Venus*, *UASp-Venus-Fim-ABD2*, and *U6-Fim-gRNA*. The *attP2* site was used for all transgenic insertions.

For RNA interference and targeted protein degradation, crosses were carried out at 18°C and embryos were collected at 18°C. For all other experiments, crosses were carried out at 25°C and embryos were collected at room temperature or 25°C.

*RNA interference*
To generate Myo-II KD embryos (Figs. 1D-1F and S1G-S1I), the progeny of *mat$\alpha$-tubulin67/UASp-zipper-shRNA; mat$\alpha$-tubulin15/+* were analyzed and control embryos were the progeny of *mat$\alpha$-tubulin67/+; mat$\alpha$-tubulin15/+*. To analyze Canoe localization in Myo-II KD embryos (Figs. S1B-S1E), the progeny of *mat$\alpha$-tubulin67/UASp-zipper-shRNA; Canoe-YFP/+* were analyzed and control embryos were the progeny of *mat$\alpha$-tubulin67/+; Canoe-YFP/+*. To analyze Fimbrin-YFP localization in *canoe* KD embryos (Fig. S4A), the progeny of Fimbrin-YFP/Fimbrin-YFP; *UAS-canoe-shRNA/mat$\alpha$-tubulin67, sqh-sqh-mCherry* were analyzed and control embryos were the progeny of *Fimbrin-YFP; mat$\alpha$-tubulin67, sqh-sqh-mCherry/+*.

*Analysis of Fimbrin localization*
To analyze Fimbrin localization in live embryos, the progeny of *Fimbrin-YFP/Fimbrin-YFP or Y; mat$\alpha$-tubulin15, sqh-gap43-mCherry/+* (Figs. 2A-2C), *Fimbrin-YFP/Fimbrin-YFP or Y; sqh-MoesinABD-mCherry/+* (Figs. 2B and S2B), *Fimbrin-YFP/Fimbrin-YFP* or *Y; sqh-sqh-mCherry/sqh-sqh-mCherry* (Figs. 2D-2J and S2C-S2I), or *Fimbrin-YFP/Fimbrin-YFP or Y; shg-shg-mCherry/+* (Figs. 2K and 2L) were analyzed.

*Analysis of cell adhesion, contractility, and behavior in Fimbrin mutants*
To analyze F-actin, Canoe, and E-cadherin localization in fixed embryos (Figs. 3A-C, 3F, 3G, 4A-4C, S3C, S4B, S4D, S6A, and S6B), the progeny of *Fimbrin$^{6}$/Fimbrin$^{6}$* or Y were analyzed and *y,w* was the wild-type control. To analyze peak retraction velocity in *Fimbrin* mutants (Figs. 5K and S3F), the progeny of *Fimbrin$^{6}$/Fimbrin$^{6}$ or Y; arm-arm-GFP/+* were analyzed and control embryos were the progeny of *Sco/CyO; arm-arm-GFP/TM6b*. To analyze Ajuba-GFP localization in *Fimbrin* mutants (Fig. S4C), the progeny of *Fimbrin$^{6}$/Fimbrin$^{6}$* or Y; *Ajuba-GFP/Ajuba-GFP* were analyzed and control embryos were the progeny of *Ajuba-GFP/Ajuba-GFP*. To analyze Canoe localization in live *Fimbrin* mutant embryos (Figs. 3J-3L), the progeny of *Fimbrin$^{6}$/Fimbrin$^{6}$* or Y; Canoe-YFP/+ were analyzed and control embryos were the progeny of *Sp/+; Canoe-YFP/+*. To analyze myosin-II localization and turnover in *Fimbrin* mutants (Figs. 5E-5H and S8F), the progeny of *Fimbrin$^{6}$/Fimbrin$^{6}$* or Y*; sqh-sqh-GFP/sqh-sqh-GFP* were analyzed and control embryos were the progeny of *y, w*; ; *sqh-sqh-GFP/sqh-sqh-GFP*. To analyze other aspects of myosin-II localization in *Fimbrin* mutants (5A, 5B, 5I, 5J, S3D and S3E), the progeny of *Fimbrin$^{6}$/Fimbrin$^{6}$* or Y*; Spider-GFP, sqh-sqh-mCherrry/+* were analyzed and control embryos were the progeny of *Sp/+; Spider-GFP, sqh-sqh-mCherry/+*. To analyze cell behavior in *Fimbrin* mutants (Figs. 4E-4G, 4I, S6H, S6I, and S7A-S7D), the progeny of *Fimbrin$^{6}$/Fimbrin$^{6}$* or Y*; Spider-GFP/Spider-GFP* were analyzed and control embryos were the progeny of *Spider-GFP/Spider-GFP*.

*Targeted degradation of Fimbrin-YFP*
For nanobody-mediated Fimbrin protein degradation in fixed (Figs. S4E, S4F and S6C-S6G) and live (Figs. S4H, S8E, S8I, S8L, S8N, and S8O) Fim KD embryos, the progeny of *Fimbrin-YFP/Fimbrin-YFP; mat$\alpha$-tubulin67, sqh-sqh-mCherry/UASp-Nslmb-VHH-GFP* were analyzed and control embryos were the progeny of *Fimbrin-YFP/Fimbrin-YFP; mat$\alpha$-tubulin67, sqh-sqh-mCherry/+*. Embryos with >70% depletion of Fimbrin-YFP levels were included in the analysis.

*Generation of Fimbrin$^{K588R}$ embryos*
To generate Fimbrin$^{K588R}$ embryos, males bearing a *UASp-Fim$^{K588R}$-Venus* or *UASp-Fim$^{K588R}$-mCherry* transgene were crossed to *mat$\alpha$-tubulin15* females bearing additional transgenes for different experiments detailed below. As maternal expression of wild-type Fimbrin or Fimbrin$^{K588R}$ significantly reduced egg laying, all analyses with Fimbrin$^{K588R}$ were performed in live embryos. To analyze myosin-II-GFP turnover in Fimbrin$^{K588R}$ embryos (Figs. 5E, 5F, 5H, and S8G), the

progeny of *sqh-sqh-GFP/+; mat$\alpha$-tubulin15/UASp-Fim*$^{K588R}$*-mCherry* were analyzed and control embryos were the progeny of sqh-sqh-GFP/+; *mat$\alpha$-tubulin15*/+. To analyze cell behavior, retraction velocity, and embryonic lethality in Fimbrin$^{K588R}$ embryos (Figs. 4E, 4F, 4H, 4J, 5K, S6H, S7F-S7J, and S8P), the progeny of *arm-arm-GFP/+; mat$\alpha$-tubulin15/UASp-Fim*$^{K588R}$*-mCherry/+* were analyzed and the control embryos were the progeny of *arm-arm-GFP/+; mat$\alpha$-tubulin15/+*. To analyze myosin-II localization in Fimbrin$^{K588R}$ embryos (Figs. 5C, 5D, 5G, S8C, S8D, S8J, and S8M), the progeny of *mat$\alpha$-tubulin15, sqh-sqh-mCherry/UASp-Fim*$^{K588R}$*-Venus* were analyzed and control embryos were the progeny of *mat$\alpha$-tubulin15, sqh-sqh-mCherry/+*. To analyze F-actin span in Fimbrin$^{K588R}$ embryos (Figs. 3D and 3E), the progeny of *sqh-Utrophin-mCherry/+; mat$\alpha$-tubulin15/UASp-Fim*$^{K588R}$*-Venus* were analyzed and control embryos were the progeny of *sqh-Utrophin-mCherry/+; mat$\alpha$-tubulin15 /+*. To analyze Canoe localization in Fimbrin$^{K588R}$ embryos (Figs. 3H and 3I), the progeny of *mat$\alpha$-tubulin15, Canoe-YFP/UASp-Fim*$^{K588R}$*-mCherry* were analyzed and control embryos were the progeny of *mat$\alpha$-tubulin15, Canoe-YFP /+*.

*Analysis of other actin crosslinkers*
To analyze Fascin localization, the progeny of *UAS-GFP-Fascin*; *mat$\alpha$-tubulin15, sqh-sqh-mCherry/+* were analyzed. The localization of other actin crosslinkers was analyzed using the *$\beta_H$-spectrin-YFP* (*kst*$^{CPTI002266}$), and *Filamin-YFP* (*cheerio*$^{CPTI001399}$) endogenous TRAP lines obtained from the Kyoto *Drosophila* Stock Center (*6*). $\alpha$-actinin localization was analyzed using an endogenous GFP tag (*3*).

**Generation of the *Fimbrin*$^{6}$ allele**
The *Fimbrin*$^{6}$ null allele was generated by targeted CRISPR-mediated deletion using published methods (*18*). Two gRNAs targeting *Fimbrin* exons 2 and 5 spanning a 1.6 kb region of the *Fimbrin* gene were cloned into the *pCFD4* plasmid (*18*) (see Plasmid construction section) and the *pCFD4-U6-Fim-gRNA* plasmid was inserted into attP2 using phiC31-mediated recombination. Females expressing *nanos-Cas9* (attP2) were crossed to males bearing *pCFD4-U6-Fim-gRNA* and F1 females expressing Cas9 and the gRNAs were crossed to *FM7/Y* males. Single F2 females were crossed to *FM7/Y* males to generate stable lines and deletions were identified by PCR screening and sequencing. Homozygous and hemizygous *Fimbrin*$^{6}$ flies were recovered and used for further analysis. Flies lacking Fimbrin are semi-viable, producing very few offspring. The reduced viability of *Fimbrin*$^{6}$ mutant embryos was rescued by supplying wild-type *Fimbrin* on BAC CH322-74M02 (*20*) inserted in attP2 (Fig. S3B).

**Immunofluorescence**
Embryos were dechorionated for 1 min in 50% bleach, washed using deionized water for 2 min, fixed for 14 min in a 1:1 mixture of heptane and 18.5% formaldehyde (Sigma) in PBS, and manually devitellinized in PBS. Devitellinized embryos were incubated in PBS-Triton buffer (0.1% Triton, 1% BSA in PBS) for 10 min, incubated 1 h in blocking solution (10% bovine serum albumin, 0.1% Triton in PBS), and stained overnight in antibody solution (5% bovine serum albumin, 0.1% Triton X-100 in PBS) at 4°C. To visualize F-actin with super-resolution microscopy, the fixation solution included Alexa 488-conjugated phalloidin (Thermo Fisher) at a dilution of 1:200. We found that Alexa 546- and Alexa 647-conjugated phalloidin gave poorer F-actin labeling compared to Alexa 488-conjugated phalloidin. Therefore, only Alexa 488-conjugated phalloidin was used for F-actin span analysis. Primary antibodies used were rabbit anti-Canoe (1:250) (*1*), rat anti-E-cadherin (1:50) (DSHB DCAD concentrated) (*2*), rabbit anti-GFP (1:150) (Torrey Pines), mouse anti-$\alpha$-spectrin (1:50) (DSHB), and GFP-nanobooster-488 (1:100) (Chromotek). Alexa Fluor secondary antibodies (Invitrogen) were used at 1:500 and Alexa-488-conjugated phalloidin (1:200), when used, was included with the secondary antibody incubation. For analyzing F-actin span, stained embryos were mounted in ProLong Diamond antifade mounting medium (Thermo

Fisher) and cured for 2 days prior to imaging. For all other experiments, stained embryos were mounted in ProLong Gold (Thermo Fisher) and cured for 2 days prior to imaging.

**Confocal microscopy with deconvolution**

Analysis of the F-actin span in fixed embryos (Figs. 1 and 3A-3C) was performed on a Zeiss LSM900 confocal equipped with an Airyscan2 detector and a 63X Plan Apo/1.4NA objective at 1.7X zoom to yield an effective pixel size of 42.5 nm. 10-15 images were acquired at 130 nm z-steps, channels were acquired sequentially in track mode, and 4 apical z-slices were projected for analysis. Partially processed Airy Sheppard Ring images generated in Zeiss Zen Blue software were deconvolved using the deconvolution module in Huygens Professional (Scientific Volume Imaging). For analyzing the F-actin span in live embryos (Figs. 3D and 3E), embryos were imaged on a spinning disk confocal using a Yokogawa CSU X1 scan head and a Hamamatsu OrcaFlash4.0 sCMOS camera on a Zeiss Axiovert 200 microscope equipped with a Zeiss Plan Neofluor 63X 1.4 NA objective. An additional magnification of 1.6X was applied with an optovar (Zeiss), yielding an effective pixel size of 51.9 nm. z-stacks of 21-27 z-slices were acquired at 150 nm z-steps with no camera binning and 4 z-slices in the region of the adherens junctions was projected for analysis. Automated signal-to-noise ratio and background calculations were performed separately for each channel using the Huygens Deconvolution Wizard with the default settings for all deconvolution performed.

**Confocal microscopy without deconvolution**

Imaging of Canoe (Figs. 3 and S4) and E-cadherin (Figs. 4, S4, and S6) in fixed embryos was performed on a Zeiss LSM700 confocal microscope with a Zeiss 40X Plan Apo/1.4 NA oil immersion objective. 10-15 images were acquired at 1.2 Airy units (slice thickness of 1.0 μm) using 0.49 μm z-steps with an additional optical magnification of 1.5, yielding an effective pixel size of 100 nm. A 2-3 μm region in the region of the adherens junctions was projected for analysis.

**Fluorescence recovery after photobleaching**

To analyze protein turnover (Figs. 5, S5, and S8), fluorescence recovery after photobleaching (FRAP) analysis was performed on an LSM700 scanning confocal with a Zeiss 40X Plan Apo/1.4NA oil immersion objective. Photobleaching was performed in a 25-pixel x 25-pixel region using the 488 nm laser at 100% intensity scanned for 100 iterations. Z-stacks of 1.6 μm slices were acquired at 0.75 μm z-steps. An additional optical magnification of 4.0 was applied, yielding an effective pixel size of 80 nm. A pre-bleaching image was acquired immediately before bleaching, and post-bleaching images were acquired every 2.5 s for myosin-II-GFP analysis in embryos and every 5 s for Fimbrin-Venus or Fimbrin-mCherry analysis in S2R+ cells. A maximum-intensity projection of 3 z-slices in the region of the adherens junctions (for myosin-II-GFP FRAP in embryos) or a single z-slice at the cortex (for Fimbrin-Venus FRAP in S2R+ cells) was analyzed.

**Time-lapse imaging**

For live imaging of uninjected embryos, embryos were dechorionated for 1 min in 50% bleach, washed using deionized water for 2 min, and mounted in a 1:1 mixture of halocarbon oils 27 and 700 (Sigma) between a gas-permeable membrane (YSI) and a #1.5 18 mm x 18 mm glass coverslip (Corning). For live imaging of drug-injected embryos (see Drug injections section), embryos were glued to a 24 mm x 40 mm glass coverslip (Corning) using glue made by dissolving double-sided tape (Scotch) in heptane (Sigma). Images were acquired on a spinning disk confocal with a Yokogawa CSU X1 scan head and an Excelitas PCO.Edge 4.2 sCMOS camera on a Zeiss Observer Z1 microscope equipped with a Zeiss Plan Neofluor 63X/1.4-NA oil-immersion objective or a spinning disk confocal with a Yokogawa CSU X1 scan head and a Hamamatsu OrcaFlash4.0 sCMOS camera on a Zeiss Axiovert 200 microscope equipped with Zeiss Plan Neofluor 40X 1.3

NA or Zeiss Plan NeoFluor 63X 1.4 NA oil-immersion objectives and z-stacks were acquired at 0.5 μm z-steps every 15 s for cell behavior and cross-correlation analysis. A maximum-intensity projection of containing 4-6 z-slices in the region of the adherens junctions was analyzed (see Quantification and image analysis section for marker-specific projection sizes).

**Laser ablation**
To analyze the effect of reducing tension on Fimbrin-YFP or $\beta_H$-spectrin-YFP localization at tricellular junctions in stage 7 embryos (Figs. 2I, 2J, and S2J), laser ablation of single edges was performed as described (*21*) using the Micropoint ablation system (Photonics instruments) tuned to 365 nm attached to a spinning disk confocal with a Yokogawa CSU X1 scan head and a Hamamatsu OrcaFlash4.0 sCMOS camera on a Zeiss Axiovert 200 microscope equipped with a Zeiss Plan Neofluor 63X 1.4 NA objective. Edges with clearly visible tricellular junctions were selected for ablation. Single edges were ablated using 10 single-point pulses. A pre-ablation image was acquired immediately before ablation, and post-ablation images were acquired every 5 s as 7-slice z-stacks at 0.5 μm z-steps. Ablations that damaged or bleached nearby edges or tricellular junctions were discarded. Up to 2 ablations were performed per embryo.

To analyze the effect of ectopic forces on protein localization at tricellular junctions in stage 7-8 embryos (Figs. 2K, 2L, 3J-3L, 5I, and 5J), a laser wounding assay was performed as described (*22*). Wounding was induced using the Micropoint ablation system described above. The apical domain of a single cell was damaged using 10 single-point pulses. A pre-ablation image was acquired immediately before ablation, and post-ablation images were acquired every 10 s for myosin-II-mCherry and Canoe-YFP and every 15 s for Fimbrin-YFP as 10-15 slice z-stacks at 0.5 μm z-steps. A 2x2 camera binning was applied for myosin-II-mCherry, Spider-GFP, and Fimbrin-YFP, and no binning was applied for Canoe-YFP. 4-10 z-slices in the region of the adherens junctions was projected for analysis (see Quantification and image analysis section for marker-specific projection sizes). Cells with 5-6 edges were selected for ablation. Ablations that caused large wounds that did not heal or that did not cause a detectable initial apical expansion (~50% of ablations in all genotypes, likely due to inaccurate targeting of the laser on rapidly moving cells) were discarded. A single ablation was performed per embryo.

The peak retraction velocity following edge ablation was measured in stage 7-8 embryos using the iLas ablation system (Gataca Systems) attached to a spinning disk confocal with a Yokogawa CSU X1 scan head and an Excelitas PCO.Edge 4.2 sCMOS camera on a Zeiss Observer Z1 microscope equipped with a Zeiss 40X Plan NeoFluor 1.3-NA objective. Ablations were performed using 10 pulses of 355-nm light along a 16-pixel line perpendicular to the targeted edge. A pre-ablation image was acquired immediately before ablation, and post-ablation images were acquired every 2 s as 5-slice z-stacks at 0.5 μm z-steps. The distance between the two tricellular junctions attached to the cut edge was measured immediately before and up to 8 s after ablation and the instantaneous retraction velocity was measured every 2 s. The peak velocity, which is predicted to be proportional to the pre-ablation tension at the cell interface (*23, 24*), was analyzed. Ablations that damaged or bleached nearby edges or tricellular junctions were discarded. Up to 3 ablations were performed per embryo.

**Drug injections**
Injections into the perivitelline space of stage 7-8 embryos were performed as described (*21, 25*). Dechorionated embryos were dried extensively by blotting the filter basket (Corning) with Kimwipes (Kimberly-Clark Professional) and transferred to a 4% agar block. Embryos were oriented ventral side up and glued to a 24 x 40 mm glass coverslip (Corning) using glue made by dissolving double-sided tape (Scotch) in heptane (Sigma). Embryos were dehydrated for 8-12 min

in Drierite (W.A. Hammond Company) until the perivitelline space was clearly visible. Solutions of 3 mM Rho-kinase inhibitor (Y-27632) (EMD Millipore) in water or 30 μM CalyculinA (EMD Millipore) in 30% DMSO were injected ventrally into the perivitelline space of stage 7-8 embryos. Injected solutions are predicted to be diluted 50-fold after injection. Water or 30% DMSO were injected as controls for Y-27632 and CalyculinA, respectively. Starting 2-5 min after injection, embryos were imaged on a spinning disk confocal using a Yokogawa CSU X1 scan head and a Hamamatsu OrcaFlash4.0 sCMOS camera on a Zeiss Axiovert 200 microscope equipped with a Zeiss Plan Neofluor 63X 1.4 NA objective. Images were acquired as z-stacks of 19-21 z-slices at 0.5 μm z-steps with 2x2 camera binning, yielding an effective xy pixel size of 0.166 μm. An apical region containing 4 z-slices in the region of the adherens junctions was projected for analysis.

**Cell culture and transfection**

*Drosophila* S2R+ cells (*Drosophila* Genomics Resource Center stock 150) were cultured at room temperature in Schneider's medium supplemented with 10% fetal bovine serum (F4135 Millipore Sigma). To express tagged Fimbrin variants using actin-Gal4, S2R+ cells were seeded onto 24-well culture plates and transfected using Fugene HD (Promega) following the manufacturer's protocol. The *pUASp-Fimbrin-Venus, pUASp-Fimbrin-mCherry, pUASp-Fimbrin*$^{K588R}$*-Venus*, or *pUASp-Fimbrin*$^{K588R}$*-mCherry* plasmids were co-transfected with *pAC5.1/V5-HisB-GAL4* (*26*). Plasmids were transfected at 500 ng/μl each and incubated at room temperature for 36-48 h. Transfected cells were seeded onto 35 mm dishes with a 20 mm glass bottom coverslip (CellVis D35-20-1.5H) precoated with 0.01% poly-L-lysine (Sigma) and allowed to adhere overnight prior to imaging.

**Quantification and image analysis**

*Actin span at tricellular junctions*. All analyses were performed in the ventrolateral region of stage 6-8 embryos. Analysis of the F-actin span at tricellular junctions (Figs. 1 and S3) was performed using a custom Python 3.11.8 script. For analysis of F-actin span in fixed embryos detected with phalloidin-488 (Figs. 1C-1F and 3A-3C), an inner circle of 11-pixel diameter (462 nm) and an outer circle of 25-pixel diameter (1.06 μm) centered on tricellular junctions was used to demarcate an inner circle and an outer ring around the tricellular junction in Fiji. For analysis of F-actin span in live embryos detected with Utrophin-mCherry (Figs. 3D and 3E), an inner circle of 9-pixel diameter (467 nm) and an outer circle of 21-pixel diameter (1.09 μm) centered on tricellular junctions was used to demarcate an inner circle and an outer ring around the tricellular junction in Fiji. Binarized images were generated using a threshold function that assigned the brightest 25% of pixels as positive and the number of positive pixels in the inner circle and outer ring was calculated from the binary images. All regions included pixels on their respective outer boundaries. The actin span was the ratio of the number of positive pixels in the outer ring to the number of positive pixels in the inner circle. The F-actin intensity was calculated as the mean pixel intensity in the entire circular region from the non-binarized images. Tricellular junctions with gaps in E-cadherin staining indicative of disrupted adhesion were not included in the analysis.

*Fluorescence intensity measurements.* To display fluorescence intensities (arbitrary units), all values were normalized to the maximum intensity value for all bicellular or tricellular junctions measured in a single experiment, with the maximum value set to 100. To analyze Canoe localization in Fim KD embryos, control and Fim KD embryos were mixed and stained in the same tube and distinguished by the presence or absence of Fimbrin-YFP detected with anti-GFP antibody. To analyze Canoe and E-cadherin localization in *Fimbrin* mutants, wild-type (*y, w*) embryos were pre-stained with Alexa 488-conjugated phalloidin and *Fimbrin* mutant embryos were pre-stained with Alexa 647-conjugated phalloidin before mixing. Embryos were then stained in the same tube and distinguished by the presence of the 488 or 647 fluorophores.

*Protein enrichment at tricellular junctions*. The TCJ ratio was analyzed in Fiji as described (*21*). 30-50 tricellular junctions were analyzed per embryo, with no tricellular junction sharing an edge with another tricellular junction analyzed. The mean intensity of each TCJ and the three connected BCJs was measured using the line tool in Fiji. The cytoplasmic background for each image was calculated as the average mean intensity values in 10-15 2x2 μm cytoplasmic regions and subtracted from the bicellular and tricellular junction intensities. The TCJ ratio was the ratio of the mean intensity at the tricellular junction divided by the average of the mean intensities of the three connected bicellular junctions. The TCJ ratio for Canoe was measured in fixed embryos (Figs. 3 and S4). The TCJ ratios for Canoe-YFP, Fimbrin-YFP, MoesinABD-mCherry, Gap43-mCherry, and Karst-YFP were measured in live embryos.

*Protein levels at tricellular junctions after edge ablation.* Protein localization at tricellular junctions following ablation of bicellular junctions was analyzed in Fiji. The mean intensities of the two tricellular junctions attached to the ablated edge and a single cytoplasmic region more than two cells away from the ablation site were measured immediately before ablation and at 5 s intervals after ablation. Cytoplasmic background was subtracted from the TCJ intensity at each time point and the background subtracted TCJ intensities were normalized to the pre-ablation intensity (t=0 s). As an unablated control, the intensity at tricellular junctions at least two cells away from the ablation site was measured, background subtracted, and normalized to t=0 as for the ablated tricellular junctions. Two ablated TCJs and three unablated TCJs were analyzed in each embryo.

*Protein levels at tricellular junctions after apical ablation.* Protein localization at tricellular junctions under ectopic forces following laser wounding was analyzed in Fiji using the following steps.

1. Image projection. Maximum-intensity projections of 10 z-slices in the region of the adherens junctions were generated for myosin-II-mCherry and Canoe-YFP, and maximum-intensity projections of 5 z-slices were generated for Fimbrin-YFP using the E-cadherin-mCherry channel to locate the first in-focus apical plane containing junctional signal, which was the most apical plane used for Fimbrin-YFP analysis. Bicellular and tricellular junction locations were identified using the E-cadherin-mCherry (Figs. 2K and 2L), Canoe-YFP (Figs. 3J-3L), or Spider-GFP (Figs. 5I-5J) channels.

2. Apical area and TCJ intensity measurements. The apical area of the ablated cell was measured using the freehand selection tool in Fiji. The onset of the contraction phase following apical ablation (t=0) was the time point at which the apical area of the ablated cell reached its maximum value. Apical areas were normalized to the value at t=0 and the apical area contraction rate was calculated as % change in area per min. The mean intensity at each TCJ associated with the ablated cell was measured starting at t=0. The background intensity, determined from a region outside the embryo, was subtracted from all TCJ intensity values to remove camera noise. The background-corrected TCJ intensities were normalized to the value at t=0. TCJs that contracted into higher-order vertices between t=0 and t=180 s were not included in the analysis.

3. Bleach correction. To correct for photobleaching, the mean intensity was measured at each time point in a large 66.4 μm X 66.4 μm embryo region more than two cells away from the ablation site. The background intensity from a region outside the embryo was subtracted from these measurements and the resulting values were normalized to t=0. Bleach correction was performed by dividing the normalized TCJ values in step 2 by the normalized values for the control region.

*Junction gaps.* To quantify gaps at 3- and 4-cell junctions, maximum-intensity 2 μm projections of the adherens junction region in fixed stage 6 and stage 7 embryos stained for endogenous E-cadherin were analyzed as described (*21*) and counted using the multi-point selection tool in Fiji.

*Vertex duration, edge contraction, and tissue elongation.* Vertex duration, edge contraction and tissue elongation were analyzed manually in time-lapse movies of embryos undergoing axis elongation acquired at 15 s time intervals as described (*25*). Spider-GFP and β-catenin-GFP were the cell outline markers for *Fimbrin* mutant and Fimbrin$^{K588R}$, respectively. For each movie, t=0 was the first time point when cells at the posterior end of the germband start moving posteriorly. For each movie, 30-40 edges oriented within 30° of the dorsal-ventral axis were selected and manually tracked to identify the first appearance of the 4-cell vertex and the first appearance of the new edge after vertex resolution. Vertex duration was calculated as the time between the first appearance of the 4-cell vertex and first appearance of the new edge. Edges that did not shrink to a vertex (<15% of edges in all genotypes, Fig. S6H) were not included in the analysis. To measure tissue elongation, two reference points 30-40 cell diameters apart along the anterior-posterior axis and 1-3 cell diameters away from the ventral furrow were identified at t=30 min. The distance between these reference points was measured in Fiji at t = -2.5 min, 0 min, 10 min, 20 min, and 30 min. The lengths were normalized to the length at t=0 to generate an elongation curve for each embryo (Figs. 4I and 4J).

*Fluorescence recovery after photobleaching (FRAP).* Analysis of fluorescence recovery after photobleaching was performed in Fiji using the following steps.

1. Image projection. Maximum-intensity z-projections of three 1.6-μm apical z-slices acquired at 0.75 μm z-steps were used for analysis.

2. Intensity measurements, background subtraction, and normalization. The mean intensity in the bleached region was measured immediately prior to bleaching and every 2.5-5 s after bleaching for 35-100 s. The cytoplasmic background was calculated as the lowest mean intensity of 3-5 cytoplasmic regions in a sub-apical z-plane to avoid medial myosin-II signal and was subtracted from the intensity in the bleached region at each time point. The background-subtracted intensities were normalized to the value immediately prior to bleaching.

3. Bleach correction. To correct for general photobleaching caused during post-photobleaching image acquisition, the mean intensity in a large embryo region one cell away from the bleached region was measured at each time point. Background subtraction and normalization was performed as for the bleached region. Bleach correction was performed by dividing the normalized intensity in the bleached region by the normalized embryo region intensity at each time point.

4. Calculation of the mobile fraction. The mobile fraction was calculated as the fraction of fluorescence recovered by 35 s after bleaching for myosin-II-GFP in embryos and 100 s for wild-type or K588R Fimbrin-Venus or Fimbrin-mCherry in S2R+ cells.

*Myosin-II cable formation.* Cable length and number were measured in live embryos expressing myosin-II-mCherry (for Fim KD and Fimbrin$^{K588R}$) or myosin-II-GFP (for *Fimbrin*$^{6}$) in early stage 8 embryos (at least 8 min after onset of elongation and at least 8 min before the onset of ventral divisions, which start at 20-25 min after the onset of elongation) to avoid compartment boundary cables prominent in late stage 8. A multicellular cable was defined as having a minimum of 3 vertical edges (oriented within 30° of the dorsal-ventral axis), a maximum angle of 30° between connected vertical edges, and continuous myosin-II localization along all edges of the cable.

Lengths for all cables within a 104 μm x 44 μm region at a minimum of one-cell width away from the ventral mesectoderm were measured using the segmented line tool in Fiji.

*Myosin-II cable ablation.* For ablation of multicellular cables in *Fimbrin* mutant and Fimbrin$^{K588R}$ embryos, β-catenin-GFP was the cell outline marker. For ablation of multicellular cables in Fim KD embryos, myosin-II-mCherry was the cell outline marker. The myosin-II marker used in each experiment was selected based on the chromosome location of available Gal4 drivers and the need for a cell outline marker for certain measurements. Cables were defined as having a minimum of 3 vertical edges (edges oriented within 30° of the dorsal-ventral axis) and a maximum angle of 30° between connected vertical edges. For embryos expressing myosin-II-mCherry, cables selected also displayed continuous myosin-II localization along all edges of the cable.

*Myosin-II planar polarity*. Myosin-II planar polarity was analyzed in live stage 7 embryos using Fiji. Spider-GFP was used to identify cell edges in wild-type and *Fimbrin* mutant embryos. Fimbrin$^{K588R}$-Venus was used to identify cell edges in Fimbrin$^{K588R}$ embryos. Myosin-II-mCherry was used to identify cell edges in the Fim KD and controls. A rectangular region containing 150-250 edges in the ventrolateral region, excluding the ventral furrow, was analyzed in each embryo, with all edges within the selected rectangular region analyzed. The mean intensity and angle were calculated for each edge. Cytoplasmic background was calculated by measuring mean intensity in ten 1.8 x 1.8 μm cytoplasmic regions and a single average value was calculated for each image. After background subtraction, edges were sorted by angle and planar polarity was calculated as the ratio of the average intensity of edges oriented within 30° of the anterior-posterior axis (horizontal edges) to the average intensity of edges oriented within 30° of the dorsal-ventral axis (vertical edges). Edges shorter than 4 pixels (664 nm) were not included in the analysis.

**Image processing**

Only raw, unprocessed images were used for fluorescence intensity analysis. Image processing for the purpose of display was performed in Fiji, including bleach correction and applying Look-Up Tables. For display in Figs. 3J, 5E, and 5I, individual images were corrected for photobleaching by dividing all pixel values in the image by the normalized bleach value for the respective time points from the analysis of unprocessed time-lapse movies (see Quantification and Image analysis). For display in movies, automated bleach correction was performed using the simple ratio (Movies S1-S3 and S6-S7) or histogram matching (Movies S4 and S5) method in Fiji. For Movies S1-S3, S6, and S7, images were also enlarged in Fiji with bilinear interpolation while preserving the aspect ratio.

**Image projection for automated segmentation**

Projections were generated in Napari (*27*) using custom Python scripts. Image z-stacks were acquired at each time point and automatically projected using a custom algorithm that identifies z-planes of interest in different regions of the image. Image z-stacks at each time point were separated into volumes with user-specified equal x and y dimensions and the z-plane with the maximum mean intensity (typically corresponding to the vitelline membrane) was set as the reference z-plane for each volume. We created an interactive plugin in Napari to create a composite image consisting of either (1) a maximum-intensity projection of a range of z-planes at a user-specified distance from the reference plane for each volume or (2) a maximum-intensity projection of a range of z-planes at a user-specified distance from the reference plane multiplied by the product of the pixel intensity values from the planes included in the projection. An additional contrast-enhancement step that stretched the histogram to the maximum and minimum allowable pixel values was applied to all images generated using method (2) and was optional for method (1).

### Machine learning model training and testing

For automated segmentation of cell borders in time-lapse movies, we developed a machine learning model to segment cell boundaries in movies (Zenodo DOI) using Python 3.7.6, leveraging Pytorch 1.13.1 (*28*) for machine learning specific tasks. We designed a convolutional neural network with a modified U-Net architecture commonly used for biomedical computer vision tasks (*29*). Two models were trained to segment movies of embryos expressing Spider-GFP or β-catenin-GFP as binary masks in which positive pixels denote cell boundaries and negative pixels denote the cytoplasm. The models were trained on movies of embryos that were previously segmented and corrected using SEGGA software and additional unpublished images (*30-36*). The Spider-GFP segmentation model was trained on 9,991 movie frames. The β-catenin-GFP segmentation model was trained on 1,378 movie frames. Segmentation outputs in the training dataset were converted to binary masks for model training in Python using custom MATLAB and Python scripts.

For each training dataset, equally sized, non-overlapping patches were cropped from the corresponding movie frames and binary masks, with 80% of patches used for training and 20% used for testing. During model training, the range of pixel values within each patch was normalized to values between zero and one, and image quality was randomly adjusted to increase model generalizability using several methods. To increase image sharpness, patches were sharpened by a factor of two using PyTorch (*28*) with a probability of 15%. To increase image noise, folded Gaussian distributed noise with a mean of zero and standard deviation of 0.2 was added to each patch with a probability of 25%, without exceeding a pixel value of one. Changes to image brightness, contrast, saturation, and hue were uniformly sampled between factors 0.5 and 2, 0.1 and 1.9, 0.5 and 1.5, and -0.25 and 0.25, respectively, using PyTorch (*28*). Augmented patches replaced the original patch in the data set.

To design the model, we modified the original U-Net architecture (*29*) combined with skip connections, but not deep supervision, from the U-Net++ model design (*37*). Our modified U-Net model downsampled the input image 6 times, and the number of feature channels doubled with each downsampling between 8 and 512. We also added a final convolutional layer that translates from eight feature channels to one feature channel, and a sigmoid layer to bound the output pixel values between zero and one.

To evaluate model accuracy during testing and training, we calculated training and testing loss values as the binary cross entropy between the original image and its ground truth binary mask using equation 1, where N is the number of pixels in the batch, y denotes the value of the pixel in the ground truth segmentation, and $p(y)$ denotes the value of the corresponding pixel in the predicted image. Model weights were updated using the Adam optimization algorithm implemented in PyTorch (*28*). We tested models trained on Spider-GFP and β-catenin-GFP images across a variety of learning rates, batch sizes, and image patch sizes using different hyperparameter combinations. Each model was trained until four consecutive epochs did not reduce the minimum average test loss across all images in the testing data set for a single epoch. A single model was chosen from each batch of hyperparameter searches if it had the lowest test loss in the best-performing epoch. From these, the final model used for image segmentation was selected based on the ease of manual corrections required to achieve accurate segmentation. The final Spider-GFP model had a learning rate of 0.0007, batch size of 1, and patch size of 128 pixels by 128 pixels. The final β-catenin-GFP model had a learning rate of 0.0003, batch size of 1, and patch size of 64 pixels by 64 pixels.

Equation 1:

$$-\frac{1}{N}\sum_{i=1}^{N} y_i \cdot \ln\left(p(y_i) + (1 - y_i) \cdot \ln\left(1 - p(y_i)\right)\right)$$

**Image segmentation and correction**

We used the Spider-GFP segmentation model to segment cell boundaries in time-lapse movies of wild-type and *Fimbrin* mutant embryos, and the β-catenin-GFP segmentation model to segment cell boundaries in time-lapse movies of control and $\text{Fimbrin}^{K588R}$ embryos. The output of each model is an image in which each pixel is assigned a prediction value between zero and one based on the probability that the pixel lies on a cell boundary in the image. The probability values for each pixel were binarized using a user-selected threshold for each movie based on visual inspection to minimize under- and over-segmentation). Remaining errors were automatically corrected using binary image erosion and dilation, respectively. Cells were identified in the resulting binary segmentations by finding enclosed regions of background signal using SciPy 1.11.3 (*38*) and attributing cell boundary pixels to the closest cell interior. The positions and identities of cells and their associated nodes and edges were assigned using a custom Python script. Custom MATLAB scripts with MATLAB 2020b (MathWorks) were used to import and export images into SEGGA software and region selection and manual correction were performed in SEGGA (*31*). Cell behaviors were analyzed using custom Python scripts with Python 3.11.0.

**Cell behavior analysis**

Neighbor exchanges and rosette rearrangements were identified computationally in the segmented images. Neighbor exchange events, also known as T1 processes (*39*), were identified as unique sets of cells that share a four-cell vertex, and rosettes were identified as unique sets of cells that share a vertex of five or more cells (*30*). Using these groupings, we calculated the cumulative number of T1 processes and rosettes per cell in each movie. Shrinking edges were counted if the edge was at least six pixels long at its maximum length, the edge was present for at least 45 consecutive seconds before contracting, and either the highest order vertex or another edge shared by cells in the T1 process or rosette was present for at least 2.5 consecutive minutes after the shrinking edge was last detected. Shrinking edges were scored as forming T1 processes if the edge contracted to a 4-cell vertex and did not join a higher-order vertex. Shrinking edges were scored as forming rosettes if the edge ultimately joined a higher-order vertex, either directly or after joining a 4-cell vertex. Shrinking rosette edges were limited to edges that appeared in the movie before the first four- or higher-cell vertex into which they collapse. The cumulative total of T1 processes and rosettes per cell was calculated as the total number of shrinking edges in each category that were associated with cells whose centroids were within the region of interest for at least 12.5 consecutive min after the onset of axis elongation (t=0) (referred to as tracked cells), divided by the total number of tracked cells. Edges that belonged to two tracked cells were counted twice; edges that belonged to one tracked cell were counted once.

### References

1. H. Miyamoto *et al.*, canoe encodes a novel protein containing a GLGF/DHR motif and functions with Notch and scabrous in common developmental pathways in Drosophila. *Genes Dev* **9**, 612-625 (1995).
2. H. Oda, T. Uemura, Y. Harada, Y. Iwai, M. Takeichi, A Drosophila homolog of cadherin associated with armadillo and essential for embryonic cell-cell adhesion. *Dev Biol* **165**, 716-726 (1994).
3. J. M. Lopez-Gay *et al.*, Apical stress fibers enable a scaling between cell mechanical response and area in epithelial tissue. *Science* **370**, eabb2169 (2020).
4. D. Sabino, N. H. Brown, R. Basto, Drosophila Ajuba is not an Aurora-A activator but is required to maintain Aurora-A at the centrosome. *J Cell Sci* **124**, 1156-1166 (2011).
5. B. M. McCartney *et al.*, Drosophila APC2 and Armadillo participate in tethering mitotic spindles to cortical actin. *Nature Cell Biology* **3**, 933-938 (2001).
6. C. M. Lye, H. W. Naylor, B. Sanson, Subcellular localisations of the CPTI collection of YFP-tagged proteins in Drosophila embryos. *Development* **141**, 4006-4017 (2014).
7. J. Huang, W. Zhou, W. Dong, A. M. Watson, Y. Hong, Directed, efficient, and versatile modifications of the Drosophila genome by genomic engineering. *Proc Natl Acad Sci U S A* **106**, 8284-8289 (2009).
8. A. Royou, C. Field, J. C. Sisson, W. Sullivan, R. Karess, Reassessing the role and dynamics of nonmuscle myosin II during furrow formation in early Drosophila embryos. *Mol Biol Cell* **15**, 838-850 (2004).
9. A. C. Martin, M. Kaschube, E. F. Wieschaus, Pulsed contractions of an actin-myosin network drive apical constriction. *Nature* **457**, 495-499 (2009).
10. X. Ren *et al.*, Optimized gene editing technology for Drosophila melanogaster using germ line-specific Cas9. *Proc Natl Acad Sci U S A* **110**, 19012-19017 (2013).
11. A. C. Martin, M. Gelbart, R. Fernandez-Gonzalez, M. Kaschube, E. F. Wieschaus, Integration of contractile forces during tissue invagination. *J Cell Biol* **188**, 735-749 (2010).
12. M. T. Abreu-Blanco, J. M. Verboon, S. M. Parkhurst, Cell wound repair in Drosophila occurs through three distinct phases of membrane and cytoskeletal remodeling. *Journal of Cell Biology* **193**, 455-464 (2011).
13. L. Figard, M. Wang, L. Zheng, I. Golding, A. M. Sokac, Membrane Supply and Demand Regulates F-Actin in a Cell Surface Reservoir. *Dev Cell* **37**, 267-278 (2016).
14. L. A. Perkins *et al.*, The Transgenic RNAi Project at Harvard Medical School: Resources and Validation. *Genetics* **201**, 843-852 (2015).
15. M. C. Lamb *et al.*, Fascin limits Myosin activity within Drosophila border cells to control substrate stiffness and promote migration. *Elife* **10**, e69836 (2021).
16. E. Caussinus, O. Kanca, M. Affolter, Fluorescent fusion protein knockout mediated by anti-GFP nanobody. *Nat Struct Mol Biol* **19**, 117-121 (2011).
17. D. Krueger, T. Quinkler, S. A. Mortensen, C. Sachse, S. De Renzis, Cross-linker-mediated regulation of actin network organization controls tissue morphogenesis. *J Cell Biol* **218**, 2743-2761 (2019).
18. F. Port, H. M. Chen, T. Lee, S. L. Bullock, Optimized CRISPR/Cas tools for efficient germline and somatic genome engineering in Drosophila. *Proc Natl Acad Sci U S A* **111**, E2967-2976 (2014).
19. R. M. Herrera-Perez, C. Cupo, C. Allan, A. Lin, K. E. Kasza, Using optogenetics to link myosin patterns to contractile cell behaviors during convergent extension. *Biophys J* **120**, 4214-4229 (2021).
20. K. J. Venken, Y. He, R. A. Hoskins, H. J. Bellen, P[acman]: a BAC transgenic platform for targeted insertion of large DNA fragments in D. melanogaster. *Science* **314**, 1747-1751 (2006).

21. H. H. Yu, J. A. Zallen, Abl and Canoe/Afadin mediate mechanotransduction at tricellular junctions. *Science* **370**, eaba5528 (2020).
22. L. Sheppard *et al.*, The alpha-Catenin mechanosensing M region is required for cell adhesion during tissue morphogenesis. *J Cell Biol* **222**, e202108091 (2023).
23. R. Farhadifar, J. C. Roper, B. Aigouy, S. Eaton, F. Julicher, The influence of cell mechanics, cell-cell interactions, and proliferation on epithelial packing. *Curr Biol* **17**, 2095-2104 (2007).
24. M. S. Hutson *et al.*, Forces for morphogenesis investigated with laser microsurgery and quantitative modeling. *Science* **300**, 145-149 (2003).
25. W. Razzell, M. E. Bustillo, J. A. Zallen, The force-sensitive protein Ajuba regulates cell adhesion during epithelial morphogenesis. *J Cell Biol* **217**, 3715-3730 (2018).
26. S. Simoes *et al.*, Compartmentalisation of Rho regulators directs cell invagination during tissue morphogenesis. *Development* **133**, 4257-4267 (2006).
27. J. Ahlers *et al.* (Zenodo, 2023).
28. A. Paszke *et al.*, PyTorch: An Imperative Style, High-Performance Deep Learning Library. *CoRR* abs/1912.01703, (2019).
29. O. Ronneberger, P. Fischer, T. Brox, U-Net: Convolutional Networks for Biomedical Image Segmentation. *CoRR* abs/1505.04597, (2015).
30. J. T. Blankenship, S. T. Backovic, J. S. Sanny, O. Weitz, J. A. Zallen, Multicellular rosette formation links planar cell polarity to tissue morphogenesis. *Dev Cell* **11**, 459-470 (2006).
31. D. L. Farrell, O. Weitz, M. O. Magnasco, J. A. Zallen, SEGGA: a toolset for rapid automated analysis of epithelial cell polarity and dynamics. *Development* **144**, 1725-1734 (2017).
32. A. C. Pare *et al.*, An LRR Receptor-Teneurin System Directs Planar Polarity at Compartment Boundaries. *Dev Cell* **51**, 208-221 e206 (2019).
33. A. C. Pare *et al.*, A positional Toll receptor code directs convergent extension in Drosophila. *Nature* **515**, 523-527 (2014).
34. S. Simoes, A. Mainieri, J. A. Zallen, Rho GTPase and Shroom direct planar polarized actomyosin contractility during convergent extension. *J Cell Biol* **204**, 575-589 (2014).
35. M. Tamada *et al.*, Toll receptors remodel epithelia by directing planar-polarized Src and PI3K activity. *Dev Cell* **56**, 1589-1602 e1589 (2021).
36. M. Tamada, D. L. Farrell, J. A. Zallen, Abl regulates planar polarized junctional dynamics through beta-catenin tyrosine phosphorylation. *Dev Cell* **22**, 309-319 (2012).
37. Z. Zhou, M. M. R. Siddiquee, N. Tajbakhsh, J. Liang, UNet++: A Nested U-Net Architecture for Medical Image Segmentation. *CoRR* **abs/1807.10165**, (2018).
38. P. Virtanen *et al.*, SciPy 1.0: fundamental algorithms for scientific computing in Python. *Nat Methods* **17**, 261-272 (2020).
39. D. Weaire, N. Rivier, Soap, cells and statistics—random patterns in two dimensions. *Contemporary Physics* **25**, 59-99 (1984).
